# CAS12A-TARGETED MULTIPLEXED NANOPORE SEQUENCING

**DOI:** 10.64898/2026.07.06.736710

**Authors:** Anna B. Rüegg, Robin Gehrold, Konstantinos Agathos, Sunwoo Chun, Aline Baur, Pawel Pelczar

## Abstract

Targeted long read sequencing (LRS) of native genomic DNA (gDNA) using Oxford Nanopore Technologies (ONT) is an economically and computationally accessible method for sequencing selected genomic regions without the limitations associated with amplification-based approaches. At present, efficiency, multiplexing, and scalability remain key challenges for existing targeted LRS. We have developed Cas12a-Targeted Multiplexed Nanopore Sequencing (CTM-nSeq), which combines Cas12a-targeting, DNA fragment enrichment, and optimized adapter ligation using T7 DNA ligase. Unlike previously established protocols, CTM-nSeq is compatible with the latest ONT flow cell chemistry. Performing CTM-nSeq on a single sample with an R10.4 MinION flow cell routinely yields hundreds of on-target reads. Furthermore, CTM-nSeq enables targeting of multiple loci and is the first targeted ONT sequencing method, allowing reliable, barcode-assisted multiplexing. CTM-nSeq is an efficient and accessible method for sequencing native gDNA and analysing DNA methylation, repeat expansions, and sequence integrity. As such, CTM-nSeq has a wide range of analytical and diagnostic applications.

**GRAPHICAL ABSTRACT:** 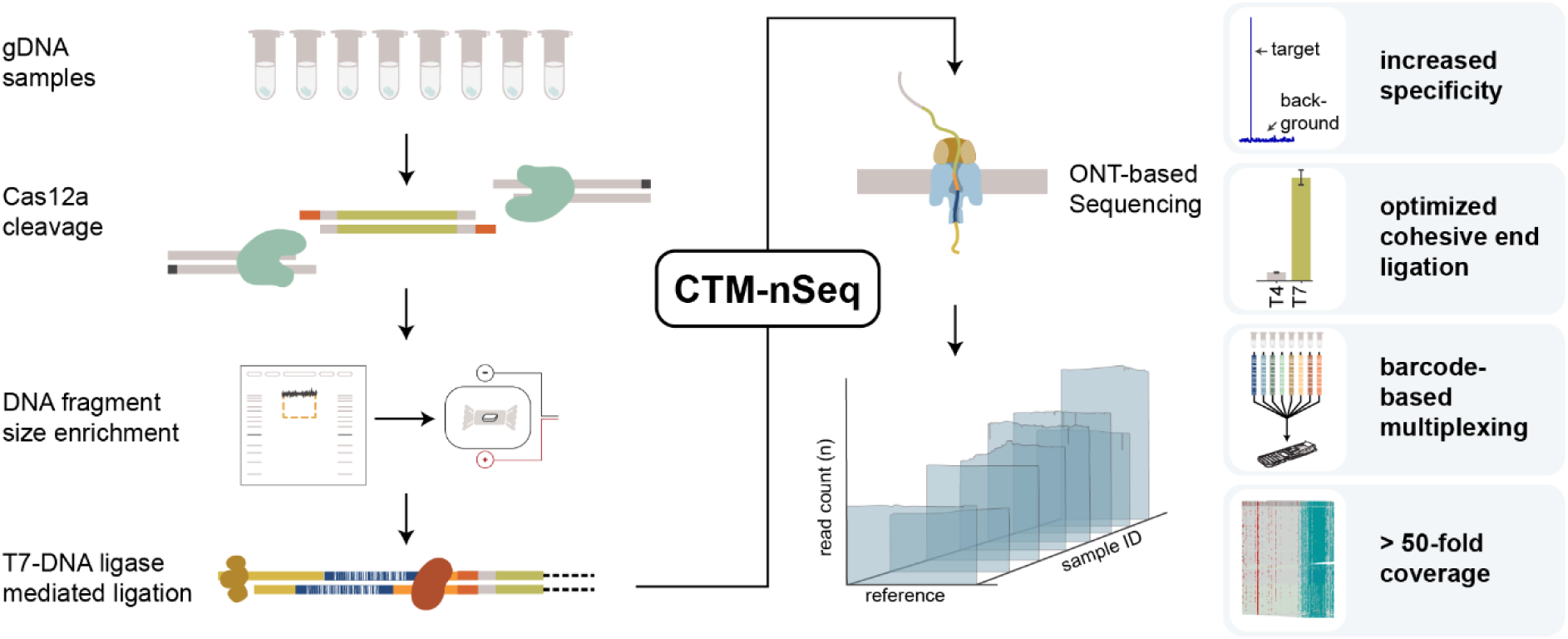

Graphical abstract illustrating CTM-nSeq workflow and advantages, showing the key steps for sample preparation: Cas12a cleavage, DNA fragment size enrichment, T7 DNA ligase mediated ligation and ONT-based sequencing. Key advantages include increased specificity, optimized cohesive end ligation, barcode-based multiplexing, and over 50-fold coverage.

## INTRODUCTION

Progress in genomics has been driven in part by long-read sequencing (LRS) technologies that permit amplification-free analysis of native genomic DNA (gDNA). LRS circumvents amplification-derived artifacts and biases, thus facilitating the analysis of previously inaccessible regions including structural variants, rearrangements, and repeats. Direct sequencing of native gDNA allows simultaneous assessment of covalent modifications, including DNA methylation. Among the LRS platforms, Oxford Nanopore Technologies (ONT) is widely used thanks to its ultralong read lengths, device portability, and low entry cost. One of the caveats of whole genome LRS is the often-prohibitive cost of achieving a genomic coverage depth sufficient for quantitative analyses at individual loci, resolution of complex sequences, or meeting standards for clinical diagnostics. In such cases, targeted nanopore sequencing offers a scalable solution that enables a deeper coverage of the selected regions. The most common targeted LRS strategies rely on PCR amplification of the target region and are, therefore, subject to PCR related size limitations, amplification bias, or failure to resolve structural rearrangements and repeats. Amplification-free targeted sequencing methods have been developed to overcome these limitations. Currently, three approaches are available namely adaptive sampling^1^, capture hybridization^2^, and Cas9-targeted enrichment (nCATS^3^). Adaptive sampling is an ONT-developed software-controlled enrichment that compares newly generated read sequences to a defined reference in real time and decides whether the sequencing read should be continued. The method can achieve up to 14-fold^1^ enrichment at target loci but is highly dependent on the correlation between predicted library fragmentation and the reference setup and requires extensive computational resources. Capture hybridization-based approaches use biotinylated capture probes combined with streptavidin magnetic bead enrichment of sequences of interest. Hybridization-based approaches are well established for short-read sequencing platforms^4–6^. However, longer sequences require prohibitively expensive probes and reports on capture hybridization in combination with LRS remain sparse^2^. nCATS leverages the sequence specificity of CRISPR/Cas9 to excise selected genomic fragments, which are subsequently ligated to ONT sequencing adapters. Dephosphorylation of gDNA prior to Cas9 cleavage enables preferential ligation of sequencing adapters to Cas9-cleavage products. nCATS has been used to validate targeted gene modifications^7,8^, study repetitive elements^9,10^, and perform targeted DNA methylation analyses^3,11^. nCATS was originally shown to achieve up to 4.6% on-target reads and a median on-target depth of up to 680 reads^3^ however, this approach does not facilitate barcode-assisted multiplexing. While nCATS performed well on R9 flow cells, adaptation to the most recent ONT R10/Kit14 chemistry, which displays improved accuracy, variation detection, and false discovery rates^13,14^, have so far remained challenging^12^.

We aimed to combine the advantages of R10/Kit14 with a new, efficient, and amplification-free targeted sequencing method that circumvents the technical challenges of nCATS. Our new targeted ONT sequencing protocol combines Cas12a-directed gDNA cleavage, DNA fragment size enrichment, and selective ligation of cohesive ends using optimized T7 DNA ligase-based adapter ligation. **C**as12-**T**argeted **M**ultiplexed **N**anopore **Seq**uencing (CTM-nSeq) is highly efficient in combination with the latest ONT chemistry, generates high numbers of on-target reads, retains the ability to probe the methylation states of native DNA, and allows barcode-based multiplexing on gDNA. As proof of principle, we applied CTM-nSeq to a selection of mouse genes and transgenic loci.

## MATERIAL AND METHODS

### Collection of tissue samples

All animals were maintained in accordance with cantonal and federal regulations and standards of the canton of Basel Stadt and Switzerland. Tissues for DNA preparation were collected from surplus animals under the license 3114_36541. The following mouse lines were used: C57BL/6JRj (wt), B6.Cg-Gt(ROSA)26Sor^tm9(CAG-tdTomato)Hze^/J (Ai9), B6.Cg-Gt(ROSA)26Sor^tm14(CAG-tdTomato)Hze^/J (Ai14), B6.Cg-Gt(ROSA)26Sor^tm32(CAG-COP4*H134R/EYFP)Hze^/J (Ai32), B6;129S-Gt(ROSA)26Sor^tm35.1(CAG-aop3/GFP)Hze^/J (Ai35D). For gDNA isolation, adult animals were euthanized using CO2. Immediately after removal, organs (spleen, pancreas, kidney, and testis) were cut into pieces of 10-20 mm3, transferred into a 1.5 ml safe-lock tube, snap frozen in liquid nitrogen, and stored at -80°C. Toe clips obtained from 1-week old pups, were collected in 1.5 ml safe-lock tubes before being frozen and stored at -20°C.

### Isolation of gDNA

HMW gDNA was isolated from tissues using the Monarch® HMW DNA Extraction Kit for Tissue (NEB, T3060) following the manufacturer’s instructions. For the isolation of HMW gDNA from toe clips, the low-input protocol of the Monarch® HMW DNA Extraction Kit for Tissue was used with the following modifications: 20 μl proteinase K in 300 μl lysis buffer for tissue lysis, only one glass bead during precipitation, and elution in 30 μl elution buffer for up to 30 minutes. For each sample, the gDNA concentration was measured using Qubit™ dsDNA Quantification Assays (ThermoFisher, Q33265/Q33230), purity was verified by spectrophotometry (A260/230 and A260/280 > 1.8), and gDNA integrity was checked by agarose gel electrophoresis.

### Design of Cas12a and Cas9 crRNAs

Cas9 Alt-R™ crRNAs (Integrated DNA Technologies, IDT) were selected using CRISPOR^47^. As Cas9 remains tightly bound to the PAM-distal site after cleavage^48^, Cas9 crRNAs were chosen such that the PAM-proximal side faces the region of interest. The position of the cleavage site and, thus, the overhang generated by Cas12a proteins have been reported to depend on the Cas12a variant and the length of the crRNA spacer^17–20,49,50^ used, with shorter spacers displaying lower overhang variability^18,20^. For this reason, we used 20 nucleotide (nt) protospacer sequences instead of the commonly used 21 nt spacer sequence. Alt-R™ A.s. Cas12a crRNA (20nt spacer, IDT) were chosen with the PAM-distal side towards the region of interest, as Cas12a was shown to remain at the PAM-proximal side^51^. Additionally, the predicted overhangs were selected to be GC-rich and screened for complementarity. All the crRNA sequences are listed in Supplementary Table 2. To facilitate the design and selection of sets with multiple crRNA pairs, a custom R-package was generated. In brief, the script uses genomic regions and DeepCpf1^21^ scores (>=30) extracted via CRISPOR^47^ as input to calculate stem-loop integrity (RNAfold^22^), specificity – no off-targets with <=1 mismatch in the first 17 bases of the protospacer (NCBI BLAST^52^), and selects for >=60% GC in the predicted overhang (nucleotides 18-23 after the PAM), to generate compatible sets of crRNAs for multiple loci such that none of the predicted overhangs have <=1 complementary base in order to avoid linker-linker duplexes.

### Design and annealing of linker and barcode oligonucleotides

One end of the linker oligonucleotides was designed to match the 4- or 5-base overhang left on the PAM-distal side between the 18^th^ and 22^nd^ or 23^rd^ nucleotide on the target strand (TS) and non-target strand (NTS), respectively (see Figure1a). The other end was designed to be complementary either directly to the NA adapter (5’-AGCAAT-3’, for simplex use) or to match part of the common sequence found in all ONT barcodes (5’-AGGTGC-3’, multiplex-linker, for use in multiplexed experiments). The centre part of the linkers was designed such that it contained a GC-clamp on each side and had a melting temperature above 50°C (5’-GGGCGAACAATGCAGG-3’). Barcode oligonucleotides were designed using the barcode sequences from ONT as a template and generating a 6-base overhang on the non-native adapter-ligated side (forward: 5’-AAGGTTAA-barcode-CAGCACC-3’, reverse: 5’-TG-barcode-TTAACCTTAGCAAT-3’). The sequences of all the linkers used are shown in Supplementary Table 4. All oligonucleotides were synthesized by Microsynth AG. Oligonucleotides were annealed in nuclease-free duplex buffer (IDT 11-01-03-01) by incubating 5 minutes at 95°C before placing the reaction on ice. Oligonucleotides were aliquoted and stored at -20°C.

**Figure 1:**
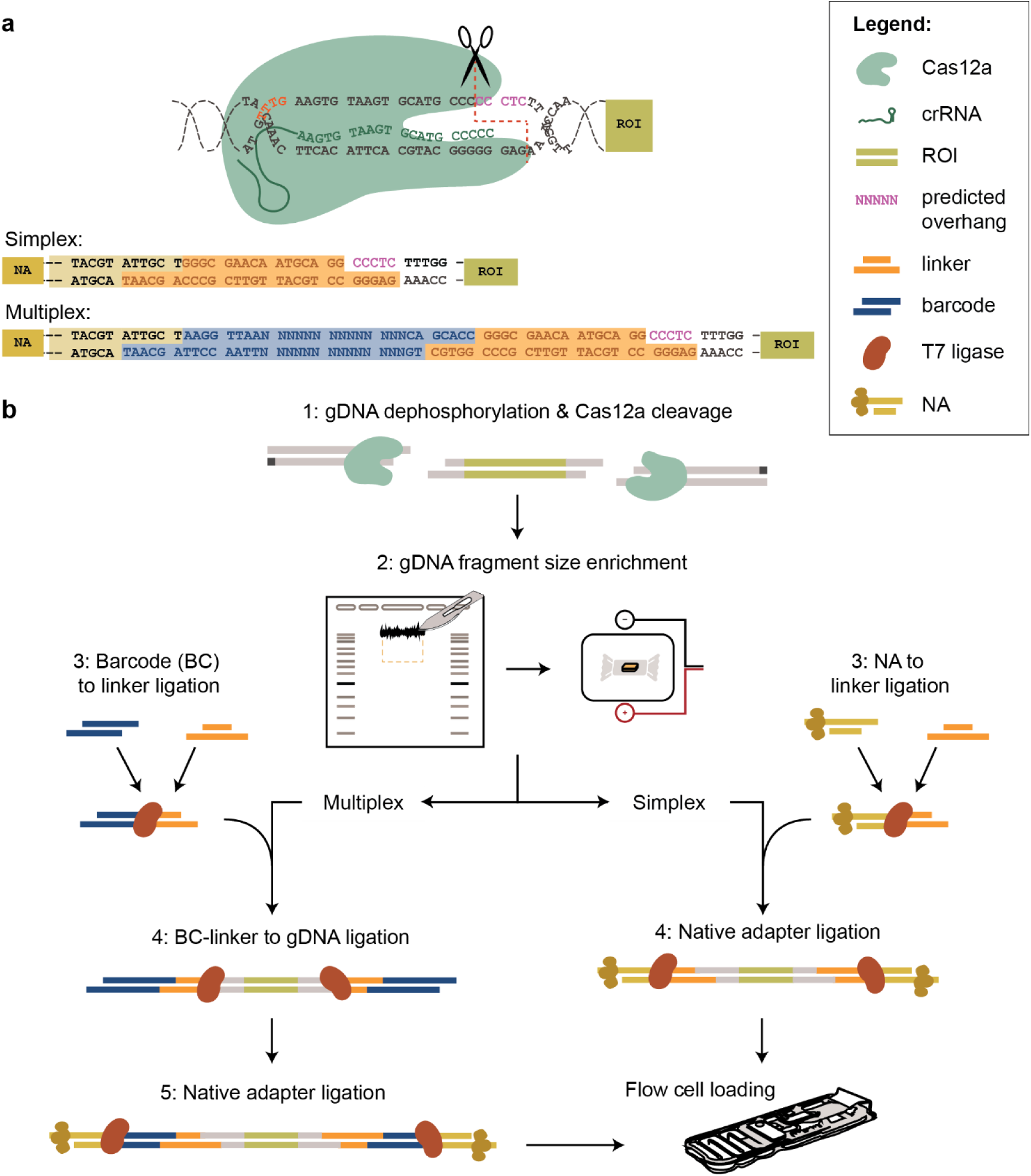
Overview of Cas12-Targeted Multiplexed Nanopore Sequencing (CTM-nSeq): a) Design of linkers and custom barcodes for simplex CTM-nSeq and multiplex CTM-nSeq: In combination with most crRNAs, AsCas12a cleaves between the 18th (non-target strand) and the 23rd (target strand) nucleotide from the PAM (TTTV), thus one end of the linker oligonucleotides is designed to match this 5-base overhang left on the PAM-distal side. The other end of the linker was designed to be complementary either directly to the ONT native adapter (NA, simplex use), or to match part of the common sequence found in all ONT barcodes (for multiplexing) b) Strategy for Cas12a enrichment: 1) gDNA is dephosphorylated and new, ligateable cohesive ends are introduced on either end of the region of interest (ROI) using AsCas12a. 2) Following Cas12a cleavage, DNA fragments are separated by agarose gel electrophoresis before selected fragments are excised and collected by electroelution into a dialysis tubing. 3) Linker oligonucleotides with overhangs complementary to the Cas12a cut sites are pre-ligated either to barcodes (BC, for multiplexing) or directly to the ONT native adapter (for simplex). 4) The linkers are subsequently ligated to the fragment size enriched gDNA. 5) For multiplexing a second round of ligation is performed to ligate the native adapter. All ligation steps are performed using T7 ligase to favour cohesive over blunt end ligation. Following preparation, the library is loaded on a ONT flow cell for sequencing.

### Ligation and analysis of small dsDNA molecules

To test the ability of T4 and T7 DNA ligases to ligate blunt, compatible cohesive, and ends with incompatible overhangs, we designed DNA oligonucleotides to generate simulated Cas12a cleavage sites, linker molecules, and NA. 10 pmoles of each oligo were incubated with either 600 U of Salt-T4® DNA Ligase (NEB, M0467S) or 900 U of T7 DNA ligase (NEB, M0318S) for 30 minutes at 25°C or using temperature cycle ligation (TCL) (30 cycles of 10 seconds at 10°C and 10 seconds at 30°C with a ramp rate of 0.5°C/s) in StickTogether™ DNA Ligase Buffer (NEB, B0535S). Samples were mixed 5:1 with purple gel loading dye (NEB, B7025S) before being loaded on a 12% Mini-PROTEAN (BioRad) native polyacrylamide gel electrophoresis (PAGE) [acrylamide/bis solution 29:1 (Bio Rad, 1610156), 0.1% w/v ammonium persulfate (Sigma, A3678), 0.05% v/v TEMED (Sigma, T9281) in 1X tris-borate-ethylenediaminetetraacetic acid (TBE) buffer (Carl Roth, 3061)]. After electrophoresis for 30-60 minutes at 4 W (Mini-PROTEAN tetra vertical electrophoresis system, Bio Rad), the gels were stained using SYBR Gold (Invitrogen, S11494) and imaged on a Gel Doc XR+ (BioRad).

### Cas12a RNP assembly

For each cleavage site, crRNAs were diluted in 1X rCutSmart buffer (NEB, B6004S) and denatured for 5 minutes at 95°C before being combined with A.s. Cas12a (Cpf1) Ultra (IDT #10001273) in 1X rCutSmart buffer at a molar ratio of 1.2 to 1 (final concentrations 2 μM Cas12a, 2.4 μM crRNA). Complexes were formed by incubating the reactions for 15 minutes at 37°C and then placed on ice or stored at 4°C for up to 24h.

### DNA cleavage and size selection

The amount of gDNA was adjusted based on the level of multiplexing (simplex on MinION flow cell: 10 μg, multiplex on MinION flow cell: 2-5 μg) and the flow cell used (simplex on Flongle flow cell: 1.3-5 μg). Prior to cleavage, gDNA was dephosphorylated in 1X rCutSmart using 3 μl QuickCIP (NEB, M0525S) per 5 μg gDNA and incubated for 20 minutes at 37°C, followed by 2 minutes at 80°C for heat inactivation. For gDNA cleavage, gDNA was mixed with ribonucleoproteins (RNPs, 2 pmol of each RNP/ μg of gDNA for up to 4 RNPs, then 0.5 pmol RNP / μg) and incubated for 60 minutes at 37°C. To inactivate Cas12, thermolabile proteinase K (0.4 μl/μg gDNA NEB, P8111S) was added and the reaction was incubated for 15 minutes at 37°C followed by 10 minutes at 55°C for heat inactivation.

### Fragment size enrichment with agarose gel electrophoresis

Following cleavage, gDNA was mixed with gel loading dye (NEB, B7024S), and run for 30 minutes at 100V on a 0.5% agarose (Bioconcept #7-01P02-R) gel in tris-acetate-ethylenediaminetetraacetic acid (TAE) buffer (Bioconcept #3-07F03-I), and stained with reduced amounts (1 μl/100 ml) of SYBR™ Safe DNA gel stain (ThermoFisher, S33102). DNA was visualized using a blue light transilluminator, and a piece of gel containing DNA fragments between 6 and 100 kb was excised and transferred into activated dialysis tubing (Spectra/Por®4, molecular weight cut-off 12-14 kD). The gDNA was electro-eluted from the gel for 30 minutes at 100V. The liquid fraction containing the gDNA was collected from the tubing and mixed with 0.4X (v/v) of Agencourt AMPure XP beads (Beckman Coulter, A63881), incubated for 10 minutes at 15 rpm at room temperature, washed twice in 80% ethanol, and eluted for 10-15 minutes at 37°C in nuclease free water.

### Linker, barcode, and adapter ligation

For simplex experiments (see Figure1b, left), 1.25 pmoles of simplex-linker and 2.5 μl of NA (ONT, EXP-NBA114) were pre-ligated in 1X quick ligation buffer for 15 minutes at 25°C using 2000 U of T7 ligase. The linker-NA ligation reaction was then mixed with cleaved and fragment size-enriched gDNA and an additional 5000 U of T7 ligase in 1X quick ligation buffer. TCL (30 cycles of 10 seconds at 10°C and 10 seconds at 30°C with a ramp rate of 0.5°C/s) was performed in 1X quick ligation, and the sample was purified using 0.3X (v/v) AMPure XP beads, washed twice with long fragment buffer (LFB, ONT, EXP-LFB001), and eluting for 10-15 minutes at 37°C in 15 μl ONT elution buffer (EB). The concentration was measured using a Qubit 1X dsDNA high sensitivity assay (ThermoFisher, Q33230), using 150-450 ng of DNA for library preparation. For the experiments performed using Flongle flow cells, the amounts were reduced by half.

For multiplex experiments (see Figure1b, right), equimolar amounts of annealed linker and barcode oligonucleotides (5 pmoles each) were pre-ligated for 15 minutes at 25°C (one reaction per barcode-linker combination) in 1X NEBNext quick ligation buffer (NEB, B6058S) using 2000 U of T7 ligase (NEB # M0318L). Pre-ligated linker-barcodes were then mixed with cleaved and fragment size-enriched gDNA and an additional 5000 U of T7 ligase in 1X NEBNext quick ligation buffer. TCL was performed as follows: 30 cycles of 10 seconds at 10°C and 10 seconds at 30°C with a ramp rate of 0.5°C/s. The ligation was stopped by adding 5% v/v ethylenediaminetetraacetic acid (EDTA) 0.5M, pH8-0 (Sigma, 324506). Samples were pooled and purified using 0.4X (v/v) AMPure XP beads, washed twice with long fragment buffer, and eluting for 10-15 minutes at 37°C in nuclease-free water. Native adapter (ONT, EXP-NBA114) ligation was performed using the TCL protocol described above by mixing the eluted, barcoded gDNA sample with 5 μl of NA and adding 10’000 U of T7 ligase in 1X NEBNext quick ligation buffer. Samples were pooled and purified using 0.3X (v/v) AMPure XP beads, washed twice with LFB, and eluted for 10-15 minutes at 37°C in 15 μl ONT elution buffer (EB). The concentration was measured using a Qubit 1X dsDNA high sensitivity assay (ThermoFisher, Q33230) and 150-450 ng of DNA was used for library preparation.

### Library preparation, flow cell loading, and ONT sequencing

Library preparation and flow cell loading were performed according to the ONT instructions for the corresponding flow cell using reagents from the Sequencing Auxiliary Vials V14 (ONT, EXP-AUX003). All experiments were performed using R10.4.1 and Kit14 chemistry and run either on a MinION flow cell (ONT, FLO-MIN114) or Flongle (ONT, FLO-FLG114). Sequencing was performed using a MinION Mk1B sequencer operated through MinKNOW (25.05.14) and allowed to run for up to 72h for MinION, 24h for Flongle flow cells.

### qPCR assays for Cas12a cleavage and ligation efficiencies

To estimate the cleavage efficiency, primers were designed to amplify a 100-200 bp fragment across the Cas12a cleavage site. An additional primer annealing to the NA sequence was designed to assess the efficiency of different ligation conditions. RNP assembly, gDNA cleavage, and PK treatment were performed as described above. After cleavage, gDNA was purified using 0.4 X Ampure XP beads before being subjected to ligation or directly used as input for qPCR. qPCR was performed using PowerUp™ SYBR™ Green Master Mix (ThermoFisher, A25742), 30-100ng (concentration measured using Qubit 1X dsDNA high sensitivity assay) of gDNA per reaction, and a primer concentration of 300nM in a final reaction volume of 10 µl using a standard protocol for the selected master mix (2 minutes 50°C, 10 minutes 95°C, 40x [15 sec 95°C, 1 minutes 60°C], ramp speed 1.6°C/s) on a QuantStudio 3 (ThermoFisher). For normalization with gDNA content, a genomic amplicon in the centre of the ROSA26 target region as well as a second target on chr12 (ApoB) were used. Sequences of all primers used are listed in Supplementary Table 3. Data was analysed using the ddCt method implemented in the “pcr” R-package^53^ and plotted using ggplot2^54^.

### Cas12a overhang characterization

To determine the overhangs generated by Cas12a after cleavage, target sites were designed as custom oligonucleotides. Each site consisted of a 17 nt sequence upstream of the PAM to introduce a BamHI overhang and a BstZ17I restriction site (5’-GATCCGAGGAGGTATAC-3’), PAM followed by 41 bases of genomic sequence and 11 additional bases to include an EcoRV cleavage site and a HindIII overhang (5’-GATATCAAGCT-3’). The target sites were cloned into BamHI-HF (NEB, R3136) and HindIII-HF (NEB, R3104) cut pBlueScript II SK+ (GeneBank X52328.1), and the insert was ligated using T4 ligase (NEB, M2200). Following plasmid cleavage with Cas12a RNP in NEB buffer 4, the reaction was inactivated by adding thermolabile proteinase K (PK, NEB, P8111) for 15 minutes at 37 °C, and the crRNA was digested using RNAse A (Invitrogen, R4642) for 10 minutes at 56 °C. For denaturing PAGE, PAM distal cleavage products were excised using EcoRV-HF (NEB, R3195) and cleaned up using sequential DNA affinity column purification (NEB, T1135, NEB, T1130). Samples were denatured in RNA Loading Dye (NEB, B0363) at 95° C for 5 minutes before being run on an 8 M Urea (Sigma 51456), 20% acrylamide (BioRad, 1610144) gel for 90 minutes at 15 W in 1x TBE buffer. Gels were stained using 2X SYBR Gold (Invitrogen, S11494) and imaged using Gel Doc XR+ (BioRad).

For ONT-based overhang determination, part of the cleaved plasmid was incubated with 1 U of Mung Bean Nuclease (MBN) (NEB, M0250) per µg of plasmid for 30 minutes at 30 °C to remove the overhang and to determine the cleavage position on the TS. To determine the cleavage position on the NTS, part of the Cas12a-cleaved plasmid was incubated with 0.5 U Q5 polymerase (Q5) (NEB, M0491) per µg of plasmid for 5 minutes at 72°C in Q5 polymerase buffer supplemented with 10 nmols dNTPs. Both reaction products were purified using a DNA affinity column (NEB, T1130) and subsequently dephosphorylated using 5 U QuickCIP (NEB, M0525S) per µg for 10 minutes at 37 °C in rCutSmart (NEB, B6004). To generate the resistance cassette, a 1048 bp fragment was amplified from pNG_v2 (Addgene #32181, forward primer: 5’-GGATTTTGGTCATGAGCTTGCGCC-3’, reverse primer: 5’-GGCAAACAAGGGGTGTTATGAGCC-3’). 150 ng of a blunt linear dsDNA fragment encoding a kanamycin resistance gene was ligated with 150 ng of processed plasmid backbone for a molar ratio of 3 parts stuffer to 1 part plasmid. The reaction used 1 µl of Quick Ligase (NEB, M2200) in Quick Ligase Buffer, supplemented with 5 mM MnCl, and incubated for 24 hours at 16 °C. The product was purified using a DNA affinity column (NEB, T1130) transformed into electrocompetent DH5α cells, and cultured on LB-kanamycin plates at 37°C for 24h. Colonies were washed off, plasmids were purified using the QIAprep Spin Miniprep Kit (Qiagen, 27104), and ONT sequenced by Microsynth AG. Data was analysed using a custom Python script that retrieved the sequences of interest using the last 40 nucleotides before the PAM and after the target site, as well as the kanamycin primers as points of reference. Putative cleavage site sequences were then aligned to the respective target site and for each position, the fraction of aligned to maximum aligned reads was calculated.

### Cas9-based enrichment including size enrichment

Cas9-based target enrichment was performed based on a combination of the protocol previously published by Gilpatrick et al.^3^ and the official ONT protocol for the native barcoding kit 24 V14 (ONT, SQK-NBD114.24 ^55^). RNP assembly was performed separately for each crRNA, Alt-R™ CRISPR-Cas9 tracrRNA (IDT 1072533) and Alt-R™ CRISPR-Cas9 crRNAs (IDT) were combined at equimolar ratios (total volume 10 μl, 10 μM each) in nuclease-free duplex buffer (IDT 11010301). The gRNA duplex was then mixed with 8 μg of Alt-R® S. pyogenes HiFi Cas9 nuclease V3, 500 μg (IDT 1081061) in 1X rCutSmart (total volume 100 μl) and incubated for 30 minutes at room temperature. In the meantime, 10 μg of gDNA was dephosphorylated with 5 μl QuickCIP (NEB, M0525S) in 1X rCutSmart, incubated for 20 minutes at 37°C, and heat inactivated for 2 minutes at 80°C. 10 μl of each Cas9 RNP, 2 μl 100mM dATP (NEB, N0440S), and 2 μl Taq DNA Polymerase (NEB, M0273L) were added to 60 μl of dephosphorylated gDNA, and the cleavage reaction was incubated for 1 hour at 37°C, followed by 10 minutes at 72°C. Agarose gel loading, excision, and product purification were performed as described for the Cas12a-based enrichment. Following gel enrichment, DNA was eluted in 20 μl nuclease free water and 12.5 μl of gDNA (about 100 ng) were mixed with 2.5 μl native barcode (ONT, SQK-NBD114.24 kit component NB) and 15 μl Blunt/TA Ligase Master Mix (NEB, M0367S). The ligation reaction was incubated for 30 minutes at 25°C before adding 6 μl 0.25M EDTA (ONT, SQK-NBD114.24 kit component) to stop the reaction and pooling the samples. Bead purification steps, adapter ligation, and library preparation were performed as described for Cas12-based enrichment.

### Data analysis

For analysis of gDNA containing ROSA26 insertions, a custom version of the mm39 reference genome was generated by inserting theoretical sequences (obtained through Addgene plasmid IDs 22799, 34880, and 34882) into the ROSA26 locus (+/- 20kb before/after homology arm) and appending all sequences to the reference genome.

All raw data was saved as .pod5 files. Raw data was basecalled using dorado 1.2.0, using hac quality presents and a minimum qscore of 10. Methylation signals were extracted using the “5mCG_5hmCG” model. Basecalled data was filtered using Filtlong v0.2.1, for a minimum read length of 500 bases, before being de-multiplexed and aligned to the mm39 reference genome or to custom references using dorado. Alignments were filtered using samtools 1.23 (using htslib 1.23) for a map quality above 20, and secondary alignments were removed. Bigwig coverage tracks were generated using bamCoverage 3.5.5. The on-target region was defined as the interval between the outermost crRNAs/gRNAs, and genome-wide and on-target sequencing statistics were calculated using samtools. Methylation statistics were calculated using modkit 0.5.0. Data was reorganized, additional statistics were calculated, and plots were generated using R version 4.5.2, ggplot2 version 4.0.2^54^, tidyr version 1.3.2^56^, and cowplot version 1.2.0^57^.

## RESULTS

### Design of crRNAs

To optimize Acidaminococcus sp. Cas12a (Cas12a) cleavage and adapter ligation conditions, we designed two pairs of CRISPR RNAs (crRNA, Supplementary Table 2) targeting the mouse safe harbour locus Gt(ROSA)26Sor (ROSA26)^15^. As Cas12a remains bound to the protospacer adjacent motif (PAM – nucleotide sequence TTTV) after cleavage^16^, we designed 20 nt crRNAs targeting the forward or reverse strand upstream or downstream of the region of interest (ROI), respectively (Figure1a). For crRNAs 20 nt or longer, cleavage predominantly occurs after the 18^th^ and 23^rd^ nucleotide from the PAM on the target strand (TS) and the non-target strand (NTS), respectively^17–20^ (Figure 1a). We selected crRNAs that meet the following criteria: DeepCpf1^21^ score > 30, limited secondary structures according to RNAfold^22^, and no off-targets with < 1 mismatch in the first 17 nt of the protospacer sequence (mm39 on NCBI BLAST). To prevent the formation of linker-dimers during subsequent ligation, we designed crRNAs such that the predicted 5-base 5’-overhangs of all cleavage products to be used in combination had not more than one complementary base-pair. In addition, we selected the predicted overhangs with a GC content > 60% to ensure efficient linker ligation. Under these constraints, crRNA design complexity increases exponentially with the number of ROIs. We created a dedicated R-package to facilitate crRNA design.

### Optimization of Cas12a cleavage, linker and adapter ligation conditions

To assess Cas12a-cleavage and linker-ligation efficiencies, we designed qPCR-based assays. We found that 2 pmoles of Cas12a were sufficient to cleave > 95% of 1 µg of gDNA in 15 minutes and 99% in 60 minutes (Supplementary Figure 1a), at which point increasing Cas12a concentration had no further effect (Supplementary Figure 1b). Given the variability of the Cas12a overhang length and position^17–20^, we assessed the ligation efficiencies of complementary linkers to putative PAM-distal overhangs. We found that linkers with both 4-base (NTS: 19, TS: 23 bases after PAM) and 5-base (NTS: 18, TS: 23 bases after PAM), but not 6-base (NTS: 18, TS: 24 bases after PAM) overhangs, were efficiently ligated to the Cas12a cut site (Supplementary Figure 1c). These results are consistent with the ONT sequencing^18^ and PAGE assays we performed to determine the PAM-distal overhangs after Cas12a cleavage (Supplementary Figure 2).

To ligate the Cas12a-cleaved gDNA to ONT’s native sequencing adapter (NA), we designed short oligonucleotide duplexes (linkers), with cohesive ends complementary to the Cas12a-generated overhang on one end and to the 6-base overhang found on ONT’s NA on the other end (Figure1a). Most adapter ligation steps in other nanopore sequencing kits rely on T/A ligation using T4 DNA ligase. However, T4 DNA ligase is less efficient at ligating longer overhangs (>4-base) and is more promiscuous in ligating incorrectly base-paired sequences compared to T7 DNA ligase^23^.

Furthermore, T7 DNA ligase is less efficient at ligating blunt ends^24^, which can decrease undesired ligation events between oligonucleotides and non-target gDNA. We confirmed that T7 DNA ligase was more efficient in ligating linkers to Cas12a-cleaved gDNA and favoured complementary cohesive ends (Supplementary Figure 1d and Supplementary Figure 3). Next, we compared the effects of ligation buffer composition, temperature, and duration on the efficiency of linker ligation to Cas12a-cleaved gDNA. We found that the fraction of ligated gDNA was higher in polyethylene glycol-containing NEBNext® quick ligation reaction buffer than in standard T4 buffer, and that the temperature cycle ligation (TCL)^25^ protocols were superior to the quick ligation protocol without requiring overnight incubation (Supplementary Figure 1e). Based on these experiments, we concluded that the combination of T7 ligase, quick ligation buffer, TCL, and a 1:1 mixture of 4- and 5-base overhang linkers was the most efficient conditions for the ligation of linkers to Cas12a-cleaved gDNA.

### DNA fragment size enrichment

In addition to optimizing the cleavage and ligation conditions, we introduced a DNA fragment size-enrichment step to mitigate over-representation of small fragments^26^ as well as to reduce extended pore occupancy and irreversible blockage caused by ultra-high-molecular-weight gDNA fragments^27^. Existing approaches to increase fragment length homogeneity use either fragmentation^27,28^, size selection^29^, or a combination of both. As none of the commercially available solutions met our needs in terms of reproducibility and adjustability, we opted for low-percentage (0.5%) agarose gel electrophoresis. The subsequent excision of fragments between ∼6 and ∼100 kb and electro-elution of the selected gDNA into dialysis tubing reduced the fraction of very short (1-6 kb) and ultra-long (>100 kb) gDNA fragments in our library preparations (Supplementary Figure 4).

### CTM-nSeq enables efficient, scalable enrichment of individual target regions

To validate CTM-nSeq, we used the Ai14^30^ mouse line, which carries a 10.6 kb reporter transgene integrated in the ROSA26 locus and targeted this ROI. We extracted, dephosphorylated, and cleaved 10 µg of spleen gDNA with two Cas12a ribonucleoproteins (RNPs) 475 bp upstream and 1420 bp downstream of the ROI. We inactivated the RNPs with thermolabile proteinase K before performing gDNA fragment size enrichment. We ligated the purified gDNA fragments directly to linker duplexes pre-ligated to ONTs NA and loaded 420 ng of the purified library onto a MinION flow cell. After base-calling and alignment with ONTs dorado, we filtered alignments with quality scores above 20 using samtools. We observed a remarkable abundance of aligned reads with an expected crRNA to crRNA read length of 12.5 kb (Figure 2a). Thus, with 582 on-target out of 8006 mapped reads (7.3%, Supplementary Figure 5, Supplementary Table 1), CTM-nSeq exceeded the previously reported on-target percentages of nCATS performed with four RNPs per ROI^3^. We achieved a mean read depth of 534 reads, highlighting that most of the reads covered the complete ROI (Figure 2b).

**Figure 2:**
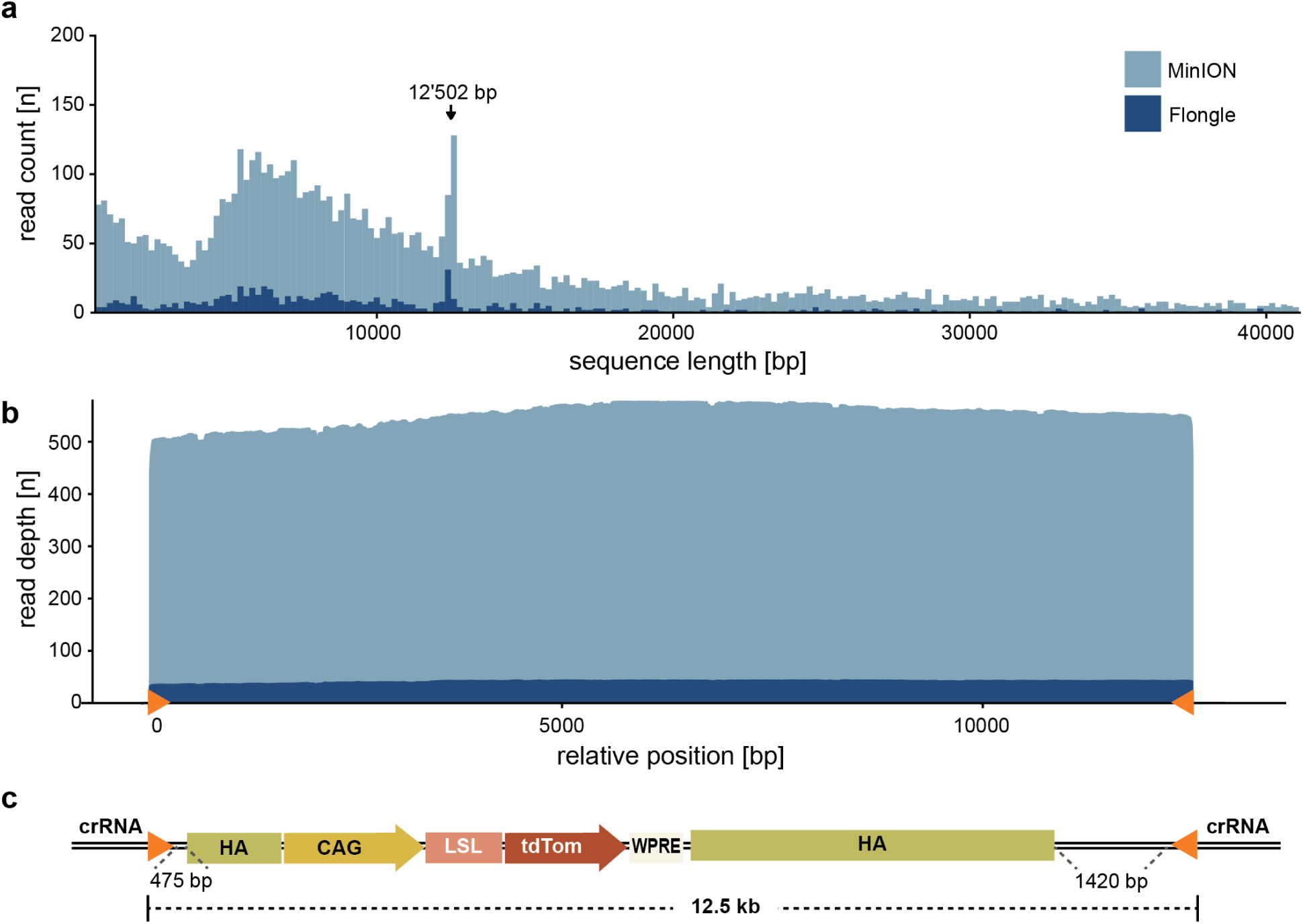
CTM-nSeq enables efficient, scalable enrichment of the target sequence using MinION and Flongle flow cells. a) Read length profile after CTM-nSeq on Ai14 mouse spleen gDNA. The library was loaded on either a MinION (light blue) or a Flongle (dark blue) flow cell. A peak corresponding to the 12.5 kb of the expected target fragment length is clearly visible (arrow). b) Read depth across the target region (crRNA to crRNA, orange triangles) achieved with a MinION (light blue) or a Flongle (dark blue) flow cell following Cas12a enrichment on Ai14 mouse spleen gDNA (the position on the x-axis is relative to the PAM of the most upstream crRNA). c) Schematic overview of Ai14 reporter transgene inserted into ROSA26 (HA: homology arms, CAG: CAG promoter, LSL: lox-stop-lox cassette, tdTom: tdTomato). The crRNAs selected for Cas12a enrichment are located 1420 and 475 bp outside of the HA, resulting in a 12’502 bp target fragment.

In parallel to sequencing on the MinION flow cell, we loaded 70 ng of the same library on a Flongle flow cell. While Flongle flow cells possess only 6-25% of the sequencing power of a MinION flow cell, the cost per flow cell is also substantially lower. As expected, the number of reads aligned to the genome, the number of on-target reads, and the median read depth were substantially lower than those achieved with a MinION flow cell (Figure 2, Supplementary Figure 5). However, the percentage of on-target reads remained comparable (5.2%; Figure 2, Supplementary Figure 5, Supplementary Table 1), and the median depth of 41 reads is still sufficient for most applications. Given that we were able to obtain a sufficiently high coverage with only a fraction (1/6) of the library as input on a Flongle flow cell, we reasoned that it might be possible to scale down the amount of input gDNA used to prepare the library. Reducing gDNA input would be of particular interest in situations where the amount of available gDNA is limited. One example is mouse transgene validation, where the biopsies are often collected as toe or ear clips. We isolated gDNA from two toe clips collected from a 10-day old mouse pup using a low-input-adapted NEB high molecular weight (HMW) gDNA extraction protocol. Starting with only 1.3 µg of gDNA for library preparation, we obtained a median depth of 7 reads (1.7% on-target reads, Supplementary Table 1) across the ROSA26 target region (wt, 7.3 kb) (Supplementary Figure 6).

### CTM-nSeq enables simultaneous enrichment of multiple target regions

A key advantage of nCATS is its ability to simultaneously target multiple loci in a single sample with limited loss of enrichment efficiency or sequencing power^3^. This allows for the creation of gene panels for repeat expansion^31,32^ or ataxia ^12^. We assessed the potential of CTM-nSeq to simultaneously target a panel of eight regions of interest: three safe-harbour loci (ROSA26, H11, and XP1) and the regulatory regions associated with five genes (Fmr1, Hnf4a, IgH, Mycbp2, and Otxr). We simplified our protocol by exclusively using 5-base overhangs, instead of a mixture of 4- and 5-base overhangs. We used 10 µg of gDNA but reduced the RNP/gDNA proportion to 0.5 pmoles/µg for each RNP to minimise the final reaction volume. Avoiding inter-linker complementarity with our crRNA design tool enabled us to perform gDNA-to-linker-NA ligation in a single reaction. We obtained 4255 on-target reads out of 33117 mapped reads (12.8%) with a median on-target read depth between 130 and 943 and an average depth of 490 (Figure 3 and Supplementary Figures 7-13, Supplementary Table 1). Interestingly, the targeted promoter region of Fmr1 was hypermethylated in approximately 50% of the reads between positions chrX:67’721’750-67’723’450. As gDNA was obtained from a female animal, this was most likely caused by X-chromosome inactivation (Supplementary Figures 7). Aligning the sequences to mm39, we observed an unexpected insertion within the ROI in IgH of approximately 3.1 kb at chr12:113’388’011 (Supplementary Figure 9-10). The sequence in question is highly repetitive and, therefore, challenging for sequencing-by-synthesis based methods. Indeed, the CTM-nSeq read sequences are 100% identical to the most recently published long-read-based genomes, C57BL_6J_T2T_v1 and C57BL_6NJ_v3^33^, indicating that the sequence found in mm39 is incomplete. This example further highlights the advantages of LRS.

**Figure 3:**
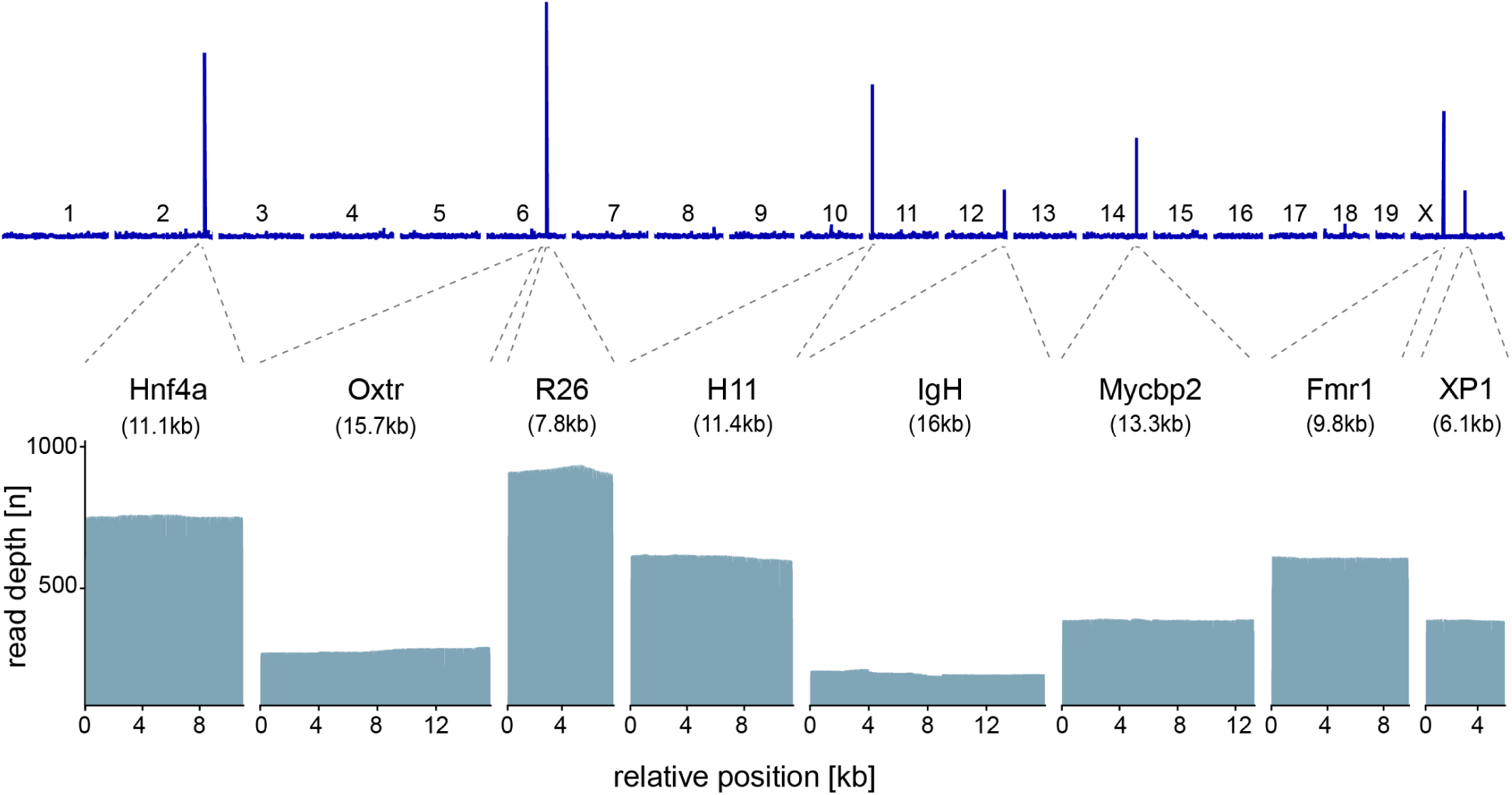
CTM-nSeq enables simultaneous enrichment of 8 genomic loci in a single reaction. The blue line in the top panel corresponds to a genome-wide coverage track, with heights showing the relative coverage for each genomic position (x-axis), and white bars separating the mm39 chromosomes indicated above. In the bottom panel, the coverage for each locus is displayed for each position between the selected crRNAs. All positions are relative to the PAM of the most upstream crRNA.

### CTM-nSeq supports barcode-based multiplexing and DNA methylation analysis

Barcoding can increase the throughput and cost-effectiveness by sequencing multiple DNA samples on a single flow cell. Existing targeted LRS methods exhibit limited barcoding compatibility. To implement barcoded CTM-nSeq, we isolated gDNA from four tissues (spleen, pancreas, kidney, and testis) of an Ai9 transgenic reporter mouse^30^ and targeted two ROIs: the ROSA26 locus and the promoter region of Hnf4a, a region known to display tissue-specific differential DNA methylation^34–36^. CTM-nSeq barcodes were designed with an NA-complementary overhang, followed by the barcode sequence, ending with a 6-base overhang complementary to a barcoding-specific version of the Cas12a cleavage site linker (Figure1a). We pre-ligated the custom barcodes with the linker molecules before ligating them to Cas12a-cleaved, fragment-enriched gDNA. After purification with magnetic beads, we performed a second ligation to add the NA and subsequently followed the standard ONT flow cell loading protocol (Figure1b). We obtained median on-target read depths between 58-275 (average 148; Figure 4a, Supplementary Table 1), with a total of 3% on-target reads (2.5-4.8% per barcode; Supplementary Table 1). We also assessed the DNA methylation profile of the Hnf4a target region. As expected, we observed promoter hypomethylation in the pancreatic gDNA – the only tissue in our sample set known to express Hnf4a^34–36^ – while the region remained hypermethylated in the spleen, testis, and kidney (Figure 4, Supplementary Figure 16). These findings highlight that DNA methylation signatures remain accessible to CTM-nSeq and sequencing depths achieved by multiplexed CTM-nSeq are suitable for differential methylation studies.

**Figure 4:**
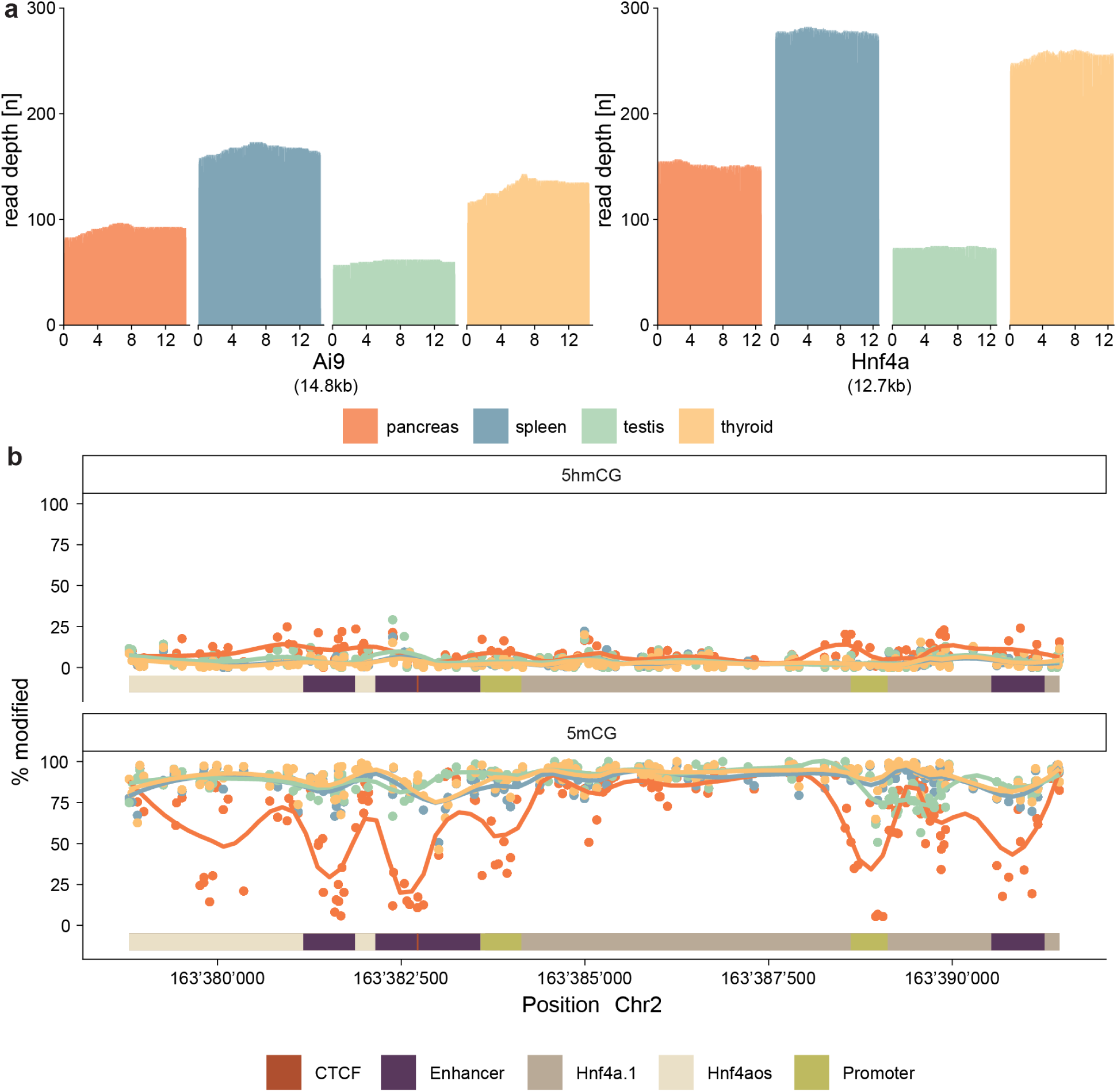
Barcode-assisted multiplexing with CTM-nSeq – simultaneous DNA sequencing and DNA methylation analyses of two ROIs in four different tissues. a) Read depth of the targeted ROIs (ROSA26 and Hnf4a) is displayed for each position between the selected crRNAs. All positions are relative to the PAM of the most upstream crRNA. b) Percentage of modified bases in the known differentially methylated targeted Hnf4a promoter region in four different tissue samples. 5mCG promoter methylation is clearly reduced in the pancreas sample which is consistent with Hnf4a being expressed in that organ, while 5hmCG levels display only minor variation.

### Optimized ligation of Cas12a-cleaved cohesive DNA ends is crucial for multiplexed targeted enrichment on R10 flow cells

Although attempts to re-establish nCATS with R10/Kit14 have been reported, the published on-target read counts and efficiencies do not meet the standards required for single nucleotide variant (SNV) calling in clinical applications^12,37^. Furthermore, nCATS with barcode-based multiplexing of mammalian gDNA has not been established for either R10 or R9 sequencing chemistry. While the cleavage- and ligation-related steps of CTM-nSeq are specific for cohesive ends generated by Cas12a, DNA fragment size selection is potentially transferable to nCATS. We adapted the nCATS protocol^38^ to include size selection and barcoding to improve the read depth and facilitate barcode-based multiplexing. We designed Cas9 crRNAs targeting sequences in close proximity to the previously used Cas12a crRNAs. We targeted the ROSA26 locus in gDNA isolated from the spleens of Ai9, Ai14, Ai32, and Ai35_D ^30,39^ (Figure 5a) mouse lines, each carrying a distinct transgene insertion in the ROSA26 locus. The modified nCATS protocol yielded 0.6 to 2. 7% on-target reads. However, the median depth was only 7-43 (Figure 5b and Supplementary Figure 17, Supplementary Table 1). In addition, 9’748 out of 16’434 reads (59%) did not contain an identifiable barcode sequence and thus remained unclassified. In the corresponding CTM-nSeq experiment, we observed comparable fractions of on-target reads (0.8 to 3.1%). Markedly, the fraction of unclassified reads was significantly lower (2’348/70’602 – 3.5%) and the median read depth was tenfold higher (140-438) compared to the modified nCATS protocol (Figure 5). We concluded that although DNA fragment size enrichment can improve the performance of nCATS with R10/Kit14, it is not sufficient to achieve clinically relevant sequencing depths. In contrast, CTM-nSeq displayed a consistent enrichment performance and generated more than the required on-target sequencing depths. We conclude that the cohesive ends generated by Cas12a and the optimised T7 DNA ligase TCL are key to increasing the specificity and efficiency.

**Figure 5:**
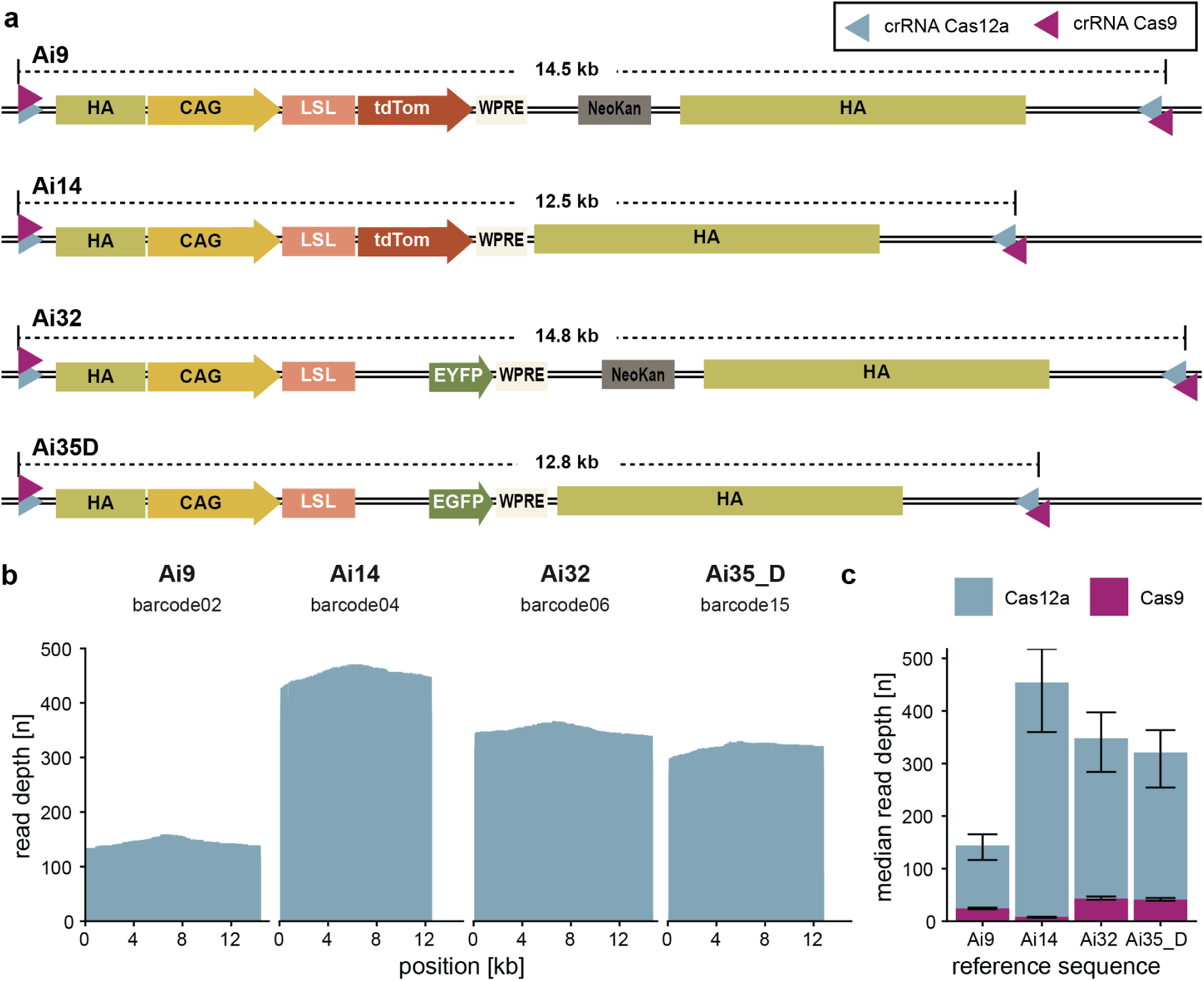
Multiplexed analysis of allelic variants in different genomic samples. a) Schematic representation of reporter alleles inserted in the ROSA26 locus (Ai9, Ai14, Ai32, Ai35_D ^30,39^). b) sequencing depths across each ROI after demultiplexing of CTM-nSeq enriched samples. All positions are relative to the PAM of the most upstream crRNA. c) Median coverage after barcoding using fragment size enrichment with size-enriched Cas9-based nCATS (purple) compared to the coverage obtained with Cas12-based CTM-nSeq (teal).

## DISCUSSION

Although the initial costs of ONT sequencing devices are low, the costs for consumables can quickly accumulate. Our aim was to establish an ONT LRS method that allows cost-efficient sequencing of multiple samples using the latest ONT chemistry. CTM-nSeq utilizes Cas12a-based cleavage, DNA fragment size enrichment, and an optimized T7 DNA ligase-based TCL protocol to achieve substantially increased on-target read counts, with several hundred end-to-end reads per target. Typical CMT-nSeq read depths are sufficient for any type of analysis, even exceeding the 50-fold coverage recommended for SNV analysis^37^.

Before the introduction of R10/Kit14, nCATS performed adequately despite its low pore occupancy^38^. The majority of DNA after nCATS is expected to be non-adapter-ligated and hence non-sequenceable, which might negatively affect flow cell performance. We found that DNA size selection also enhanced the efficiency of nCATS on R10 flow cells (Figure 5 and Supplementary Figure 17); however, modified nCATS produced roughly 10-fold fewer reads than CTM-nSeq. Size enrichment requires a substantial amount of gDNA (10 µg), and our attempts to reduce gDNA quantities resulted in limited on-target read depths (Supplementary Figure 6). Nevertheless, 10 µg of gDNA should not pose an obstacle in the clinical context, as > 30 µg/ml DNA can be isolated from human blood^40^. Furthermore, the >7-fold coverage achieved from toe biopsies-derived gDNA can be sufficient for structural and transgene integrity validation.

Perhaps the most critical element of CTM-nSeq is the generation and exploitation of cohesive DNA ends generated by Cas12a. Conversely, nCATS relies on T/A-based adapter ligation of Cas9-cleaved dA-tailed gDNA using T4 DNA ligase. Here, the nCATS approach has two disadvantages. First, sample handling may introduce new phosphorylated DNA ends by shearing, leading to unspecific ligation products. Second, the efficiency of T/A-ligation is significantly lower than that of cohesive end ligation^41^, leaving behind non-ligated target fragments. We show that substituting T4 with T7 DNA ligase reduces non-specific ligation. At the same time, Cas12a-generated 5’-overhangs in combination with a TCL protocol increase ligation efficiency (Supplementary Figure 1, Supplementary Figure 3). In line with previous observations^12^, we confirm that suboptimal ligation efficiency is a likely bottleneck for implementing barcoding with nCATS (Figure 5c).

Despite its effectiveness, the two properties of Cas12a that could limit its utility for targeted sequencing are trans-cleavage and cleavage site variability. Following on-target cleavage, the activated RuvC domain can continue to cleave or degrade single- or double-stranded RNA and DNA substrates in a crRNA-independent manner, defined as trans-cleavage^16,42–45^. Among the most frequently used Cas12a variants, AsCas12a displays the lowest trans-cleavage activity^46^. Therefore, we selected AsCas12a for CTM-nSeq. Cleavage site variability, that is, Cas12a cleavage not exclusively occurring between nucleotides 18 and 23 after the PAM^17–20^, could be a second caveat. Indeed, our overhang analysis (Supplementary Figure 4) suggests that not all generated PAM-distal overhangs are compatible with the current linker design. Nevertheless, our results suggest that a sufficient amount of the cleavage product can be ligated to adaptors with 4- and 5-base overhangs (Supplementary Figure 1). Future strategies aimed at better control over trans-cleavage activity and cleavage site variability may help to further improve the performance of CTM-nSeq.

Taken together, our results demonstrate that CTM-nSeq outperforms current targeted LRS methods, delivering superior on-target sequencing depth and enrichment. The integration of barcode-based multiplexing broadens the application spectrum and enhances the cost efficiency. As such, CTM-nSeq is a valuable tool for targeted LRS studies, such as analysing DNA methylation, repeat expansions, validating transgene integrity, and more with numerous applications for basic and applied research, as well as for clinical diagnostics.

## ACKNOWLEDGEMENTS

We thank Prof. Verdon Taylor for critical revision of the manuscript and all members of the Taylor group for discussions and input.

## AUTHOR CONTRIBUTIONS

Pawel Pelczar: Conceptualization, Methodology, Supervision, Writing—review & editing. Anna B. Rüegg: Methodology, Investigation, Formal analysis, Software, Visualization, Writing—original draft, review & editing, Robin Gehrold: Investigation, Formal analysis, Writing—review & editing. Konstaninos Agathos: Software, Formal analysis, Writing—review & editing. Sunwoo Chun and Aline Baur: Investigation, Writing—review & editing

## SUPPLEMENTARY DATA

Supplementary Data are available at NAR online.

## DATA AVAILABILITY

A step-by-step protocol is available upon request. All primary and secondary data generated has been deposited on Zenodo (10.5281/zenodo.20407579).

## SUPPLEMENTARY MATERIALS

**Supplementary Figure 1:**
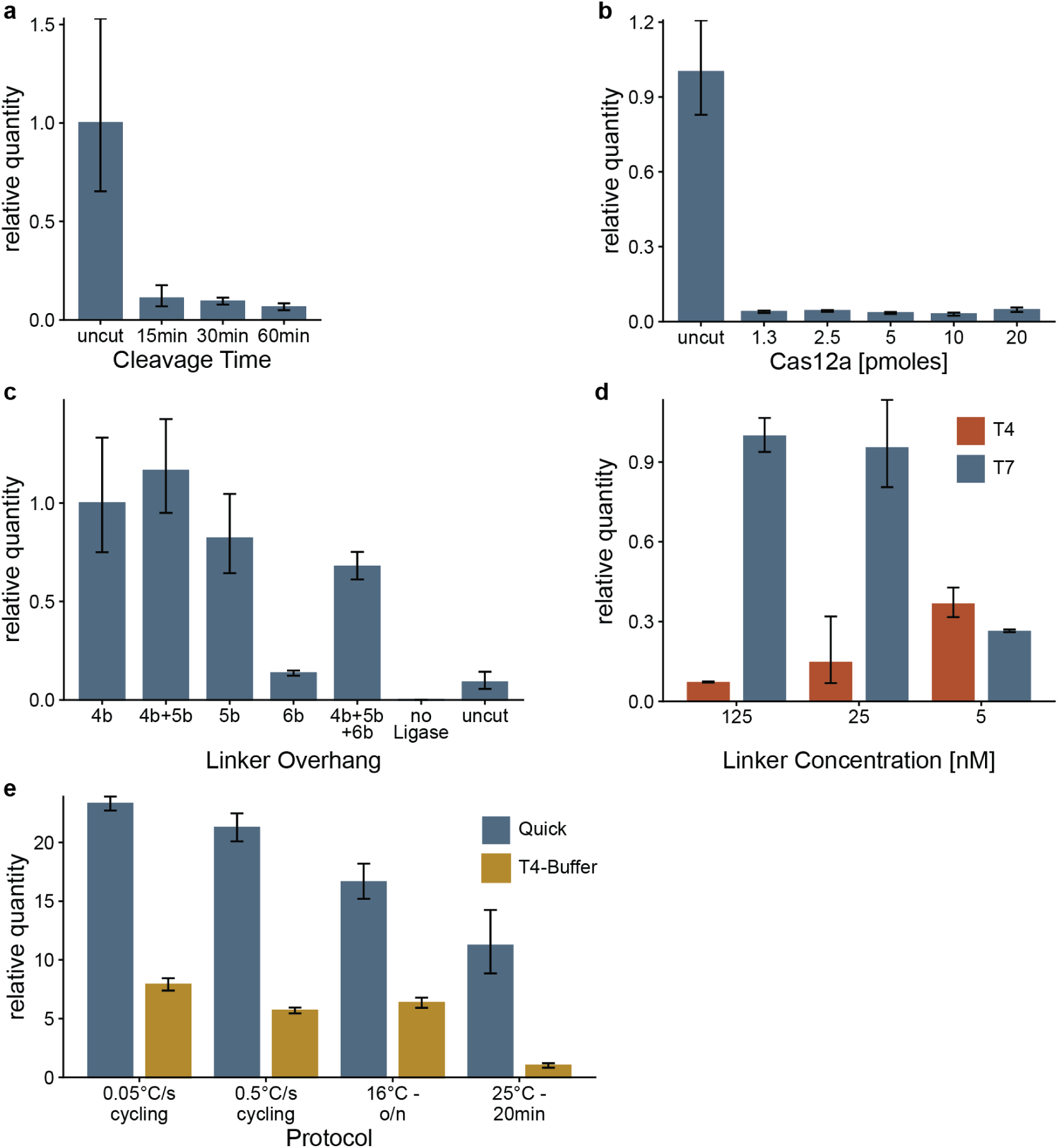
qPCR results from assays to establish CTM-nSeq reaction conditions using crRNA ROSA26_As20-440.rv. a) Time course of Cas12a cleavage reaction. All samples contained 2 pmoles of Cas12a and 1 µg of mouse gDNA). b) Titration of Cas12a concentrations in a 60-minute cleavage reaction with 1 µg of mouse gDNA. c) comparison of ligation efficiencies between Cas12a-cleaved gDNA and linkers with a variable complementary 5’-overhang. 4-base overhang (4b), 1:1 mixture of 4 & 5-base complementary overhangs (4b+5b), 5-base (5b), 6-base (6b) as well as an equimolar mixture of 4-, 5-, and 6-base overhangs (4b+5b+6b). d) Evaluation of ligation efficiency at different linker concentrations for T4 and T7 ligase. e) Evaluation of T7 DNA ligase ligation protocols: 10°C-30°C cycling at different ramping speeds, static 16 °C over-night, and 25°C 20 minute “quick” ligation in standard T4 vs NEB next quick ligation buffer.

**Supplementary Figure 2:**
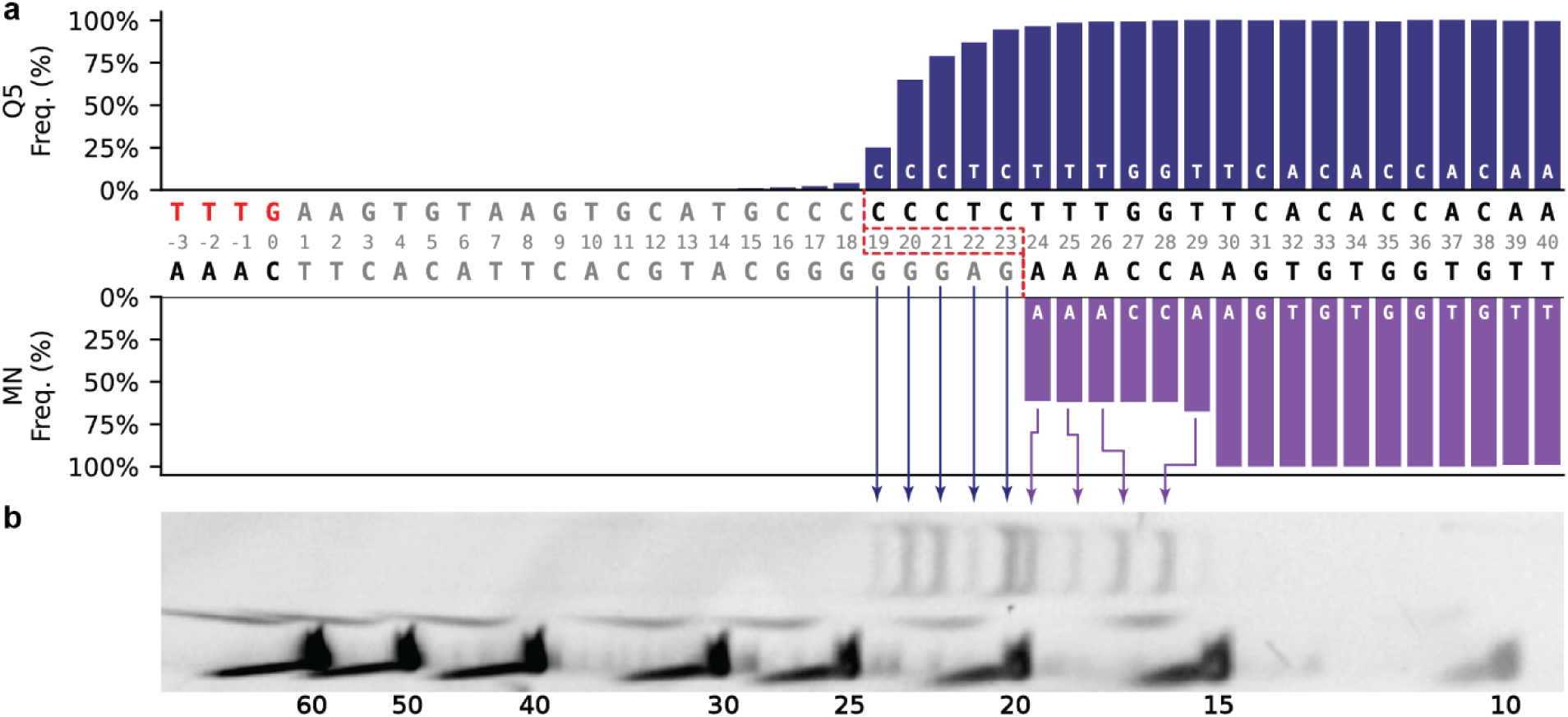
PAM distal products of cleavage with crRNA ROSA26_As20-440.rv. a) graphical representation of cleavage events observed with ONT based Cas12a cleavage site assay. The frequency was calculated as the fraction of reads aligned to each position relative to all reads aligned. Blue arrows represent Q5 processed ends and purple arrows represents MBN processed ends. Positions are relative to the PAM marked in red font. The predicted 5-base overhang is indicated with dashed red lines. b) denaturing PAGE of the same sample with lines linking the products from the two methods.

**Supplementary Figure 3:**
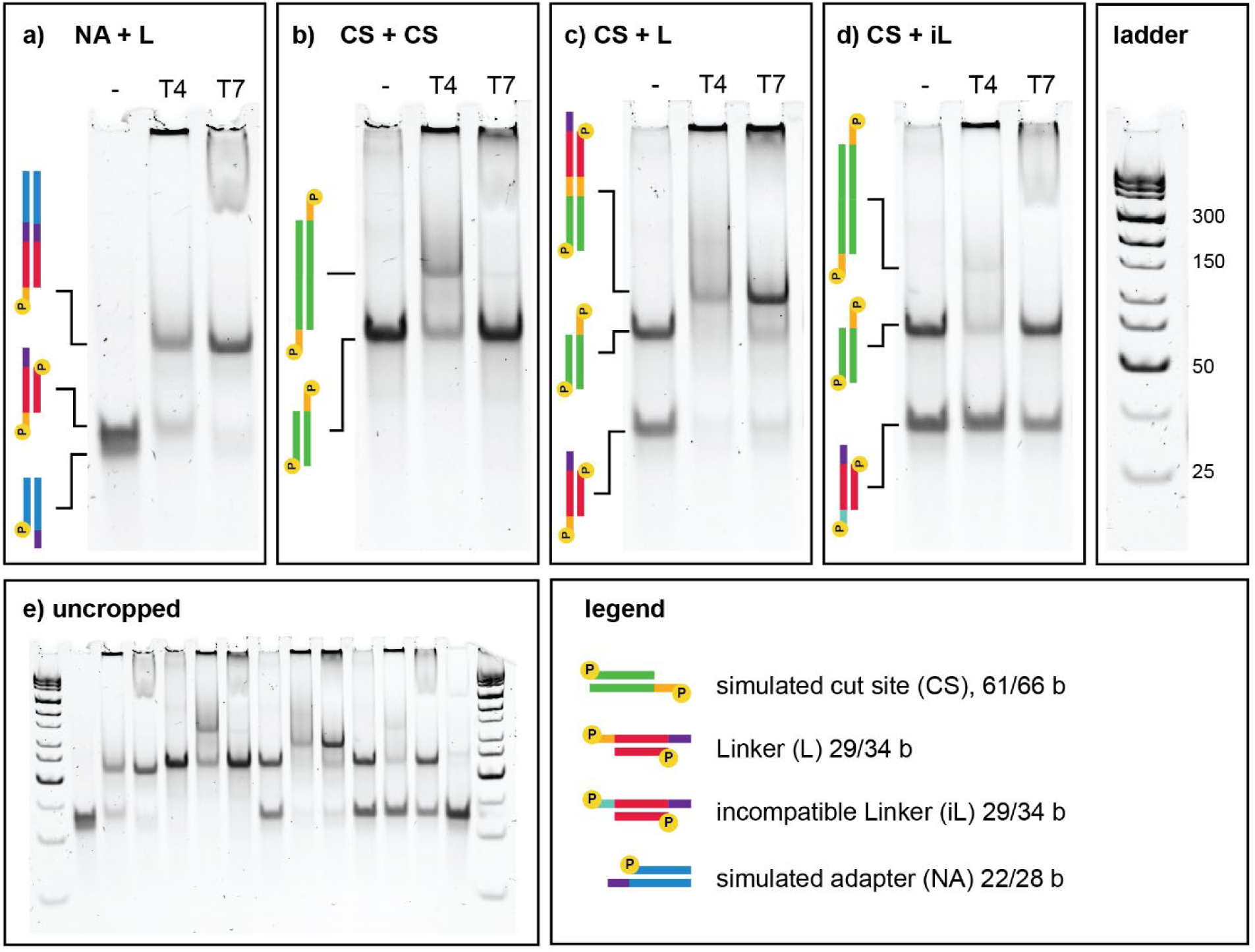
Testing the ligation of DNA blunt ends, compatible cohesive ends, and incompatible overhangs using T4 and T7 DNA ligase. Native PAGE of ligated short dsDNA fragments designed to simulate Cas12a-cleavage site, linker and adapter ligation. a) Cohesive ends are ligated more efficiently by the T7 DNA ligase when compared to the T4 DNA ligase. b) Blunt ends are ligated almost exclusively by T4 and not by T7 DNA ligase. c) presence of both cohesive and blunt ends leads to the formation of undesired long ligation products if T4 but not if T7 DNA ligase is used. d) Incompatible protruding (non-cohesive) ends are not ligated by either ligase. As seen by the DNA fragments not entering the gel but remaining in the well, T4 DNA ligase also has an increased tendency to generate oligonucleotide-multimers, probably caused by non-specific ligation. e) uncropped image of the gel shown in panels a-d.

**Supplementary Figure 4:**
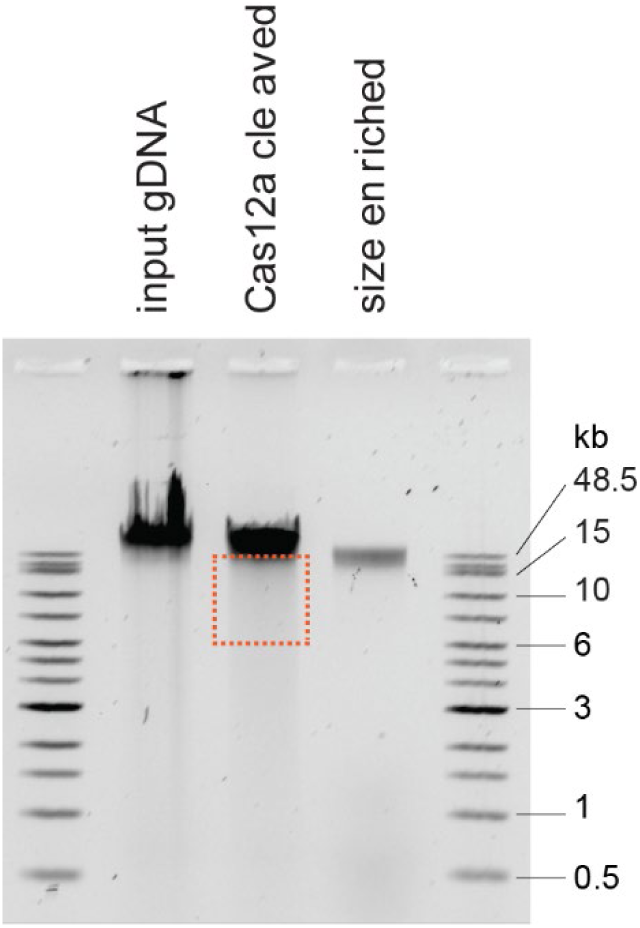
Example of a low-percentage agarose gel used for gDNA fragment size enrichment. gDNA was loaded on 0.5% TAE agarose gel after isolation (input gDNA), Cas12a cleavage and recovered fragments after gel excision and elution (size enriched). The dashed box indicates the approximate location of the excised gel fragment.

**Supplementary Figure 5:**
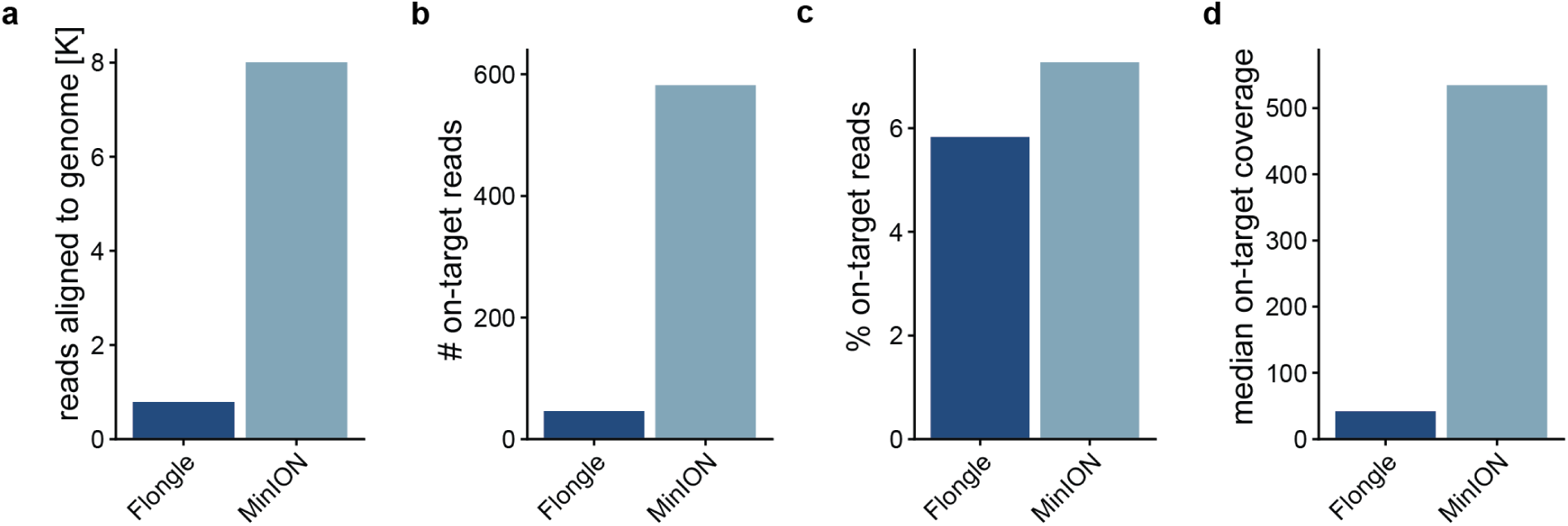
Comparison of Flongle (dark blue) and MinION (light blue) flow cell performance with respect to total number of reads aligned to the mouse genome (a), number of on-target reads (b), percentage aligned reads that align to the target region (c), and median on-target coverage (d).

**Supplementary Figure 6:**
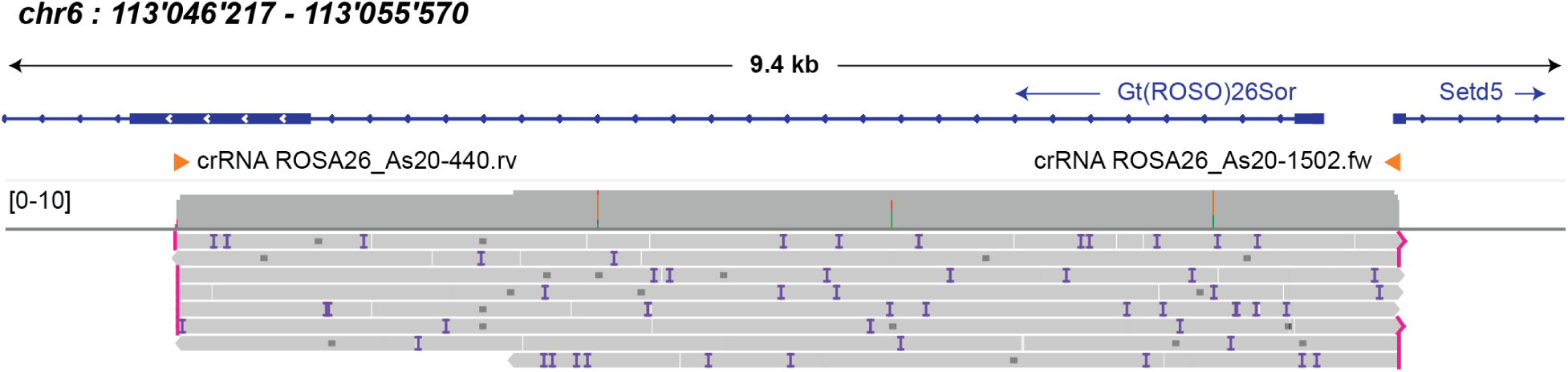
IGV snapshot of reads aligned to ROSA26 target region using gDNA form a mouse toe biopsy. gDNA was obtained from two toe clippings of the same one-week old animal before Cas12a enrichment and sequencing on a Flongle flow cell. Expanded representation allows to distinguish each read as a horizontal line. Insertions of >2 bases are shown in purple and deletions are shown as dashes. Note that most of these are related to sequencing errors as expected from ONT sequencing data.

## IGV Snapshots

**Supplementary Figures 7 to Supplementary Figure 15: ONT reads from mouse spleen gDNA aligned to genomic loci simultaneously targeted using CTM-nSeq (see also Figure 3).** Forward reads are tinted grey reverse reads light green, unmethylated (methylation probability <50%) CpGs are shown in teal, 5mC modifications are shown in red, and 5hmC modifications in orange.

**Supplementary Figures 7:**
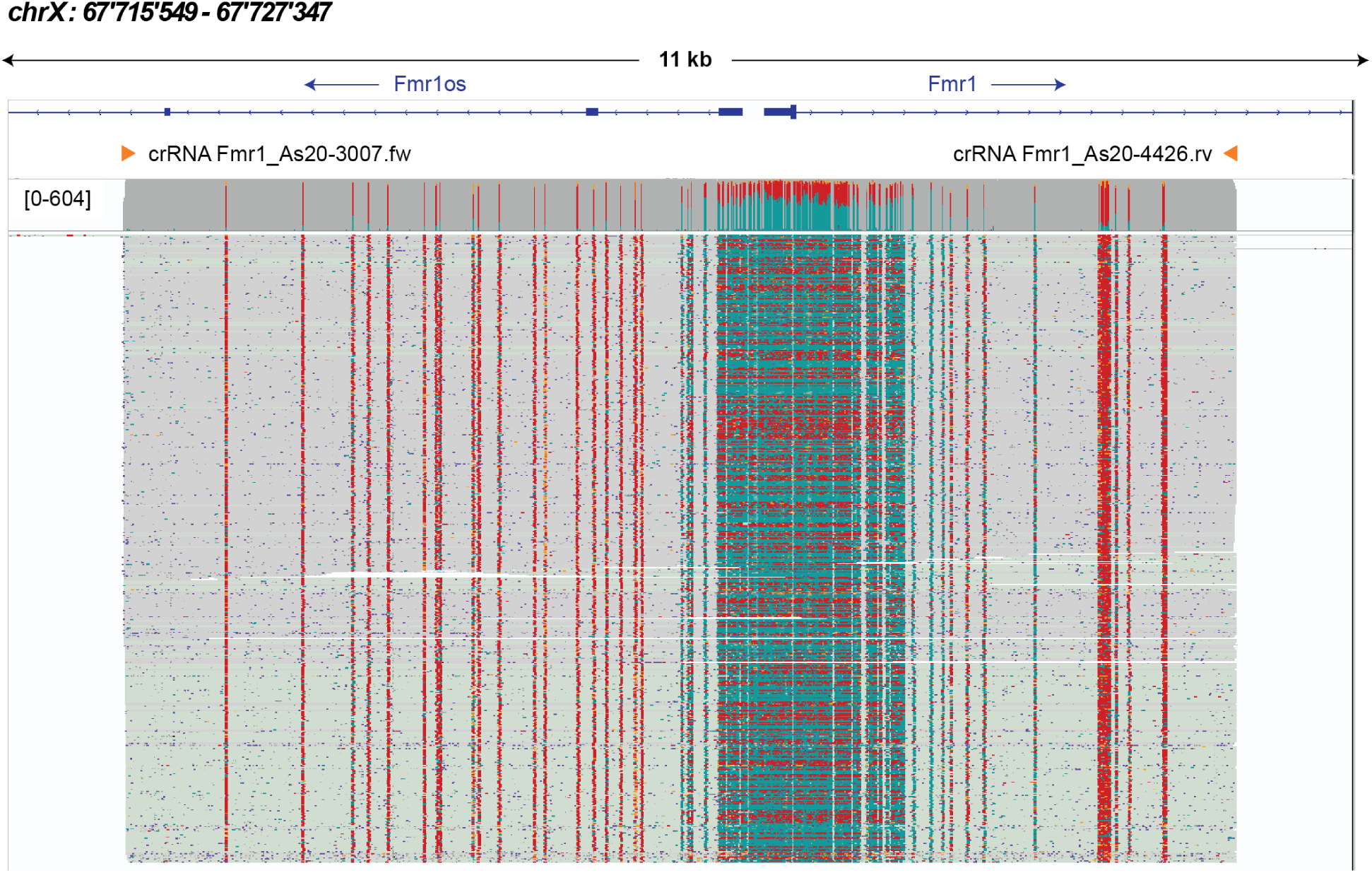
IGV-Snapshot of the reads aligned to the Fmr1 target region. Reads were obtained by simultaneously targeting 8 different loci (Supplementary Figures 7 to Supplementary Figure15, summarized in Figure 3). Forward reads are grey reverse reads light green, unmethylated (methylation probability <50%) CpGs are shown in teal, 5mC modifications are shown in red, and 5hmC modifications are shown in orange.

**Supplementary Figure 8:**
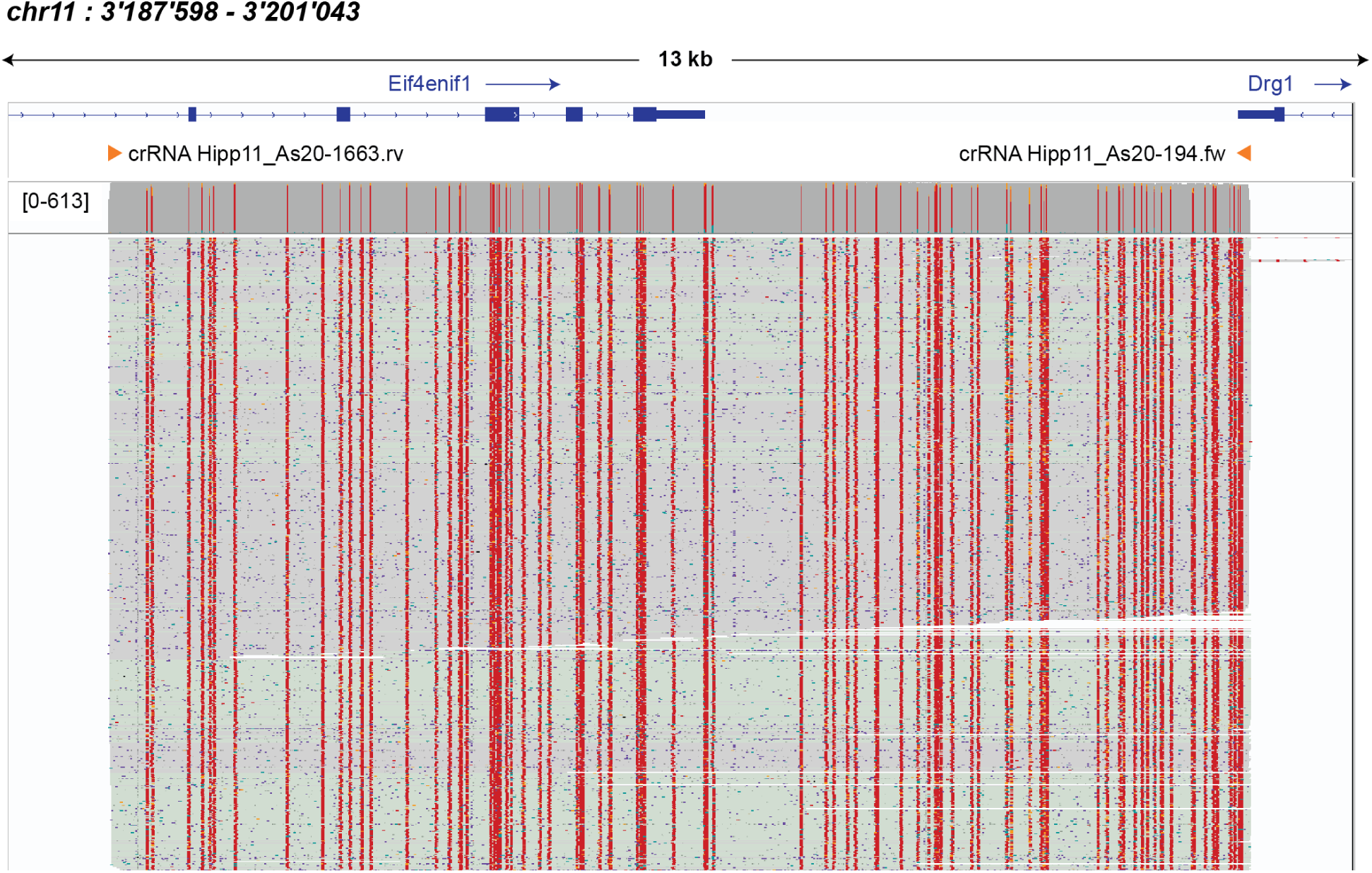
IGV-Snapshot of the reads aligned to the H11 target region. Reads were obtained by simultaneously targeting 8 different loci (Supplementary Figures 7 to Supplementary Figure15, summarized in Figure 3). Forward reads are grey reverse reads light green, unmethylated (methylation probability <50%) CpGs are shown in teal, 5mC modifications are shown in red, and 5hmC modifications are shown in orange.

**Supplementary Figure 9:**
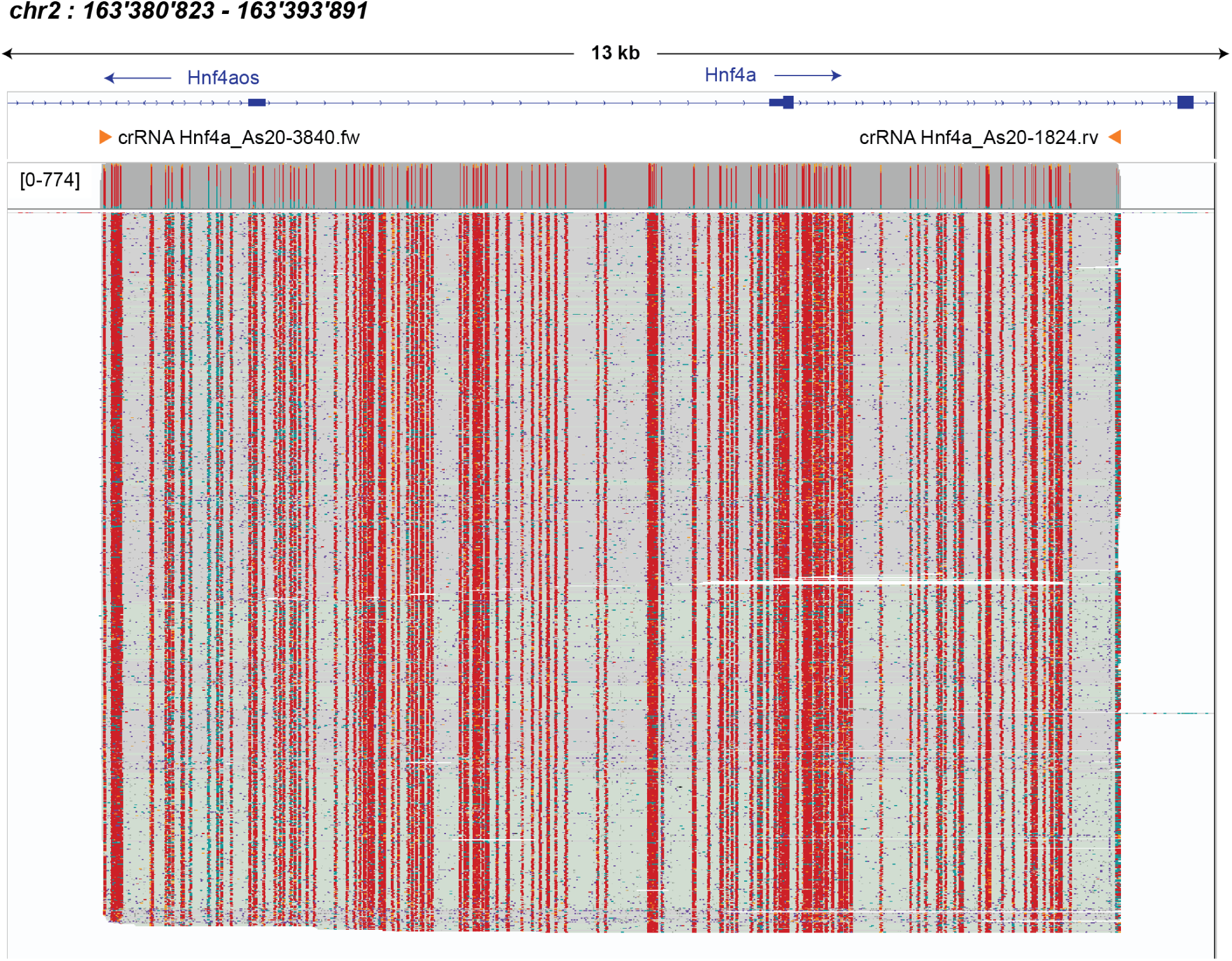
IGV-Snapshot of the reads aligned to the Hnf4a target region. Reads were obtained by simultaneously targeting 8 different loci (Supplementary Figures 7 to Supplementary Figure15, summarized in Figure 3). Forward reads are grey reverse reads light green, unmethylated (methylation probability <50%) CpGs are shown in teal, 5mC modifications are shown in red, and 5hmC modifications are shown in orange.

**Supplementary Figure 10:**
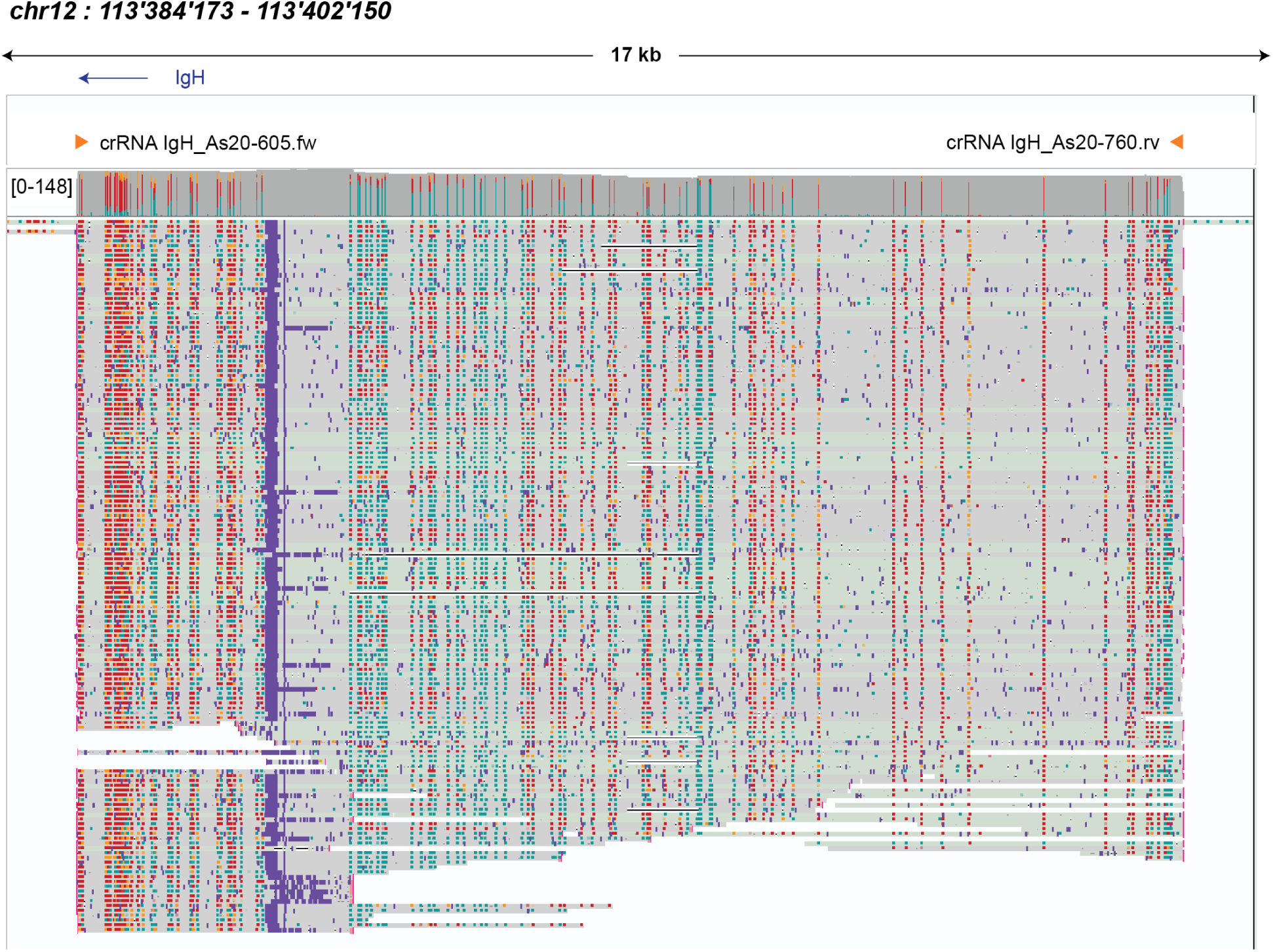
IGV-Snapshot of the reads aligned to the IgH target region. Reads were obtained by simultaneously targeting 8 different loci (Supplementary Figures 7 to Supplementary Figure15, summarized in Figure 3). Forward reads are grey reverse reads light green, unmethylated (methylation probability <50%) CpGs are shown in teal, 5mC modifications are shown in red, and 5hmC modifications are shown in orange.

**Supplementary Figure 11:**
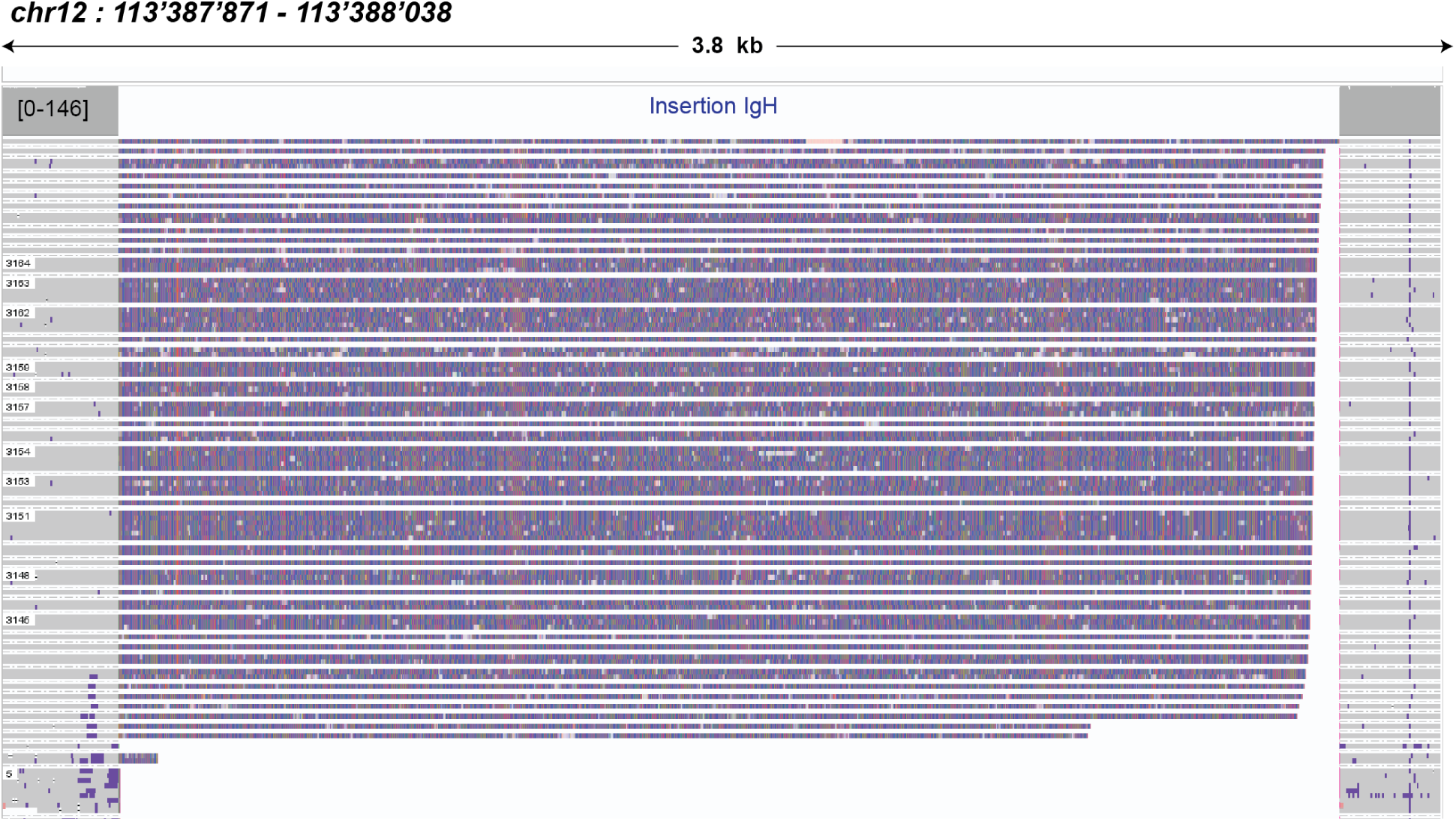
split junction track for the observed > 3 kb insertion within IgH target region. The insertion was observed when reads were aligned to the mm39 genome, but not when aligned to recent LRS genomes C57BL_6J_T2T_v1 and C57BL_6NJ_v3^33^. Therefore, the discrepancy is likely a result of the mm39 sequence being incomplete in this region.

**Supplementary Figure 12:**
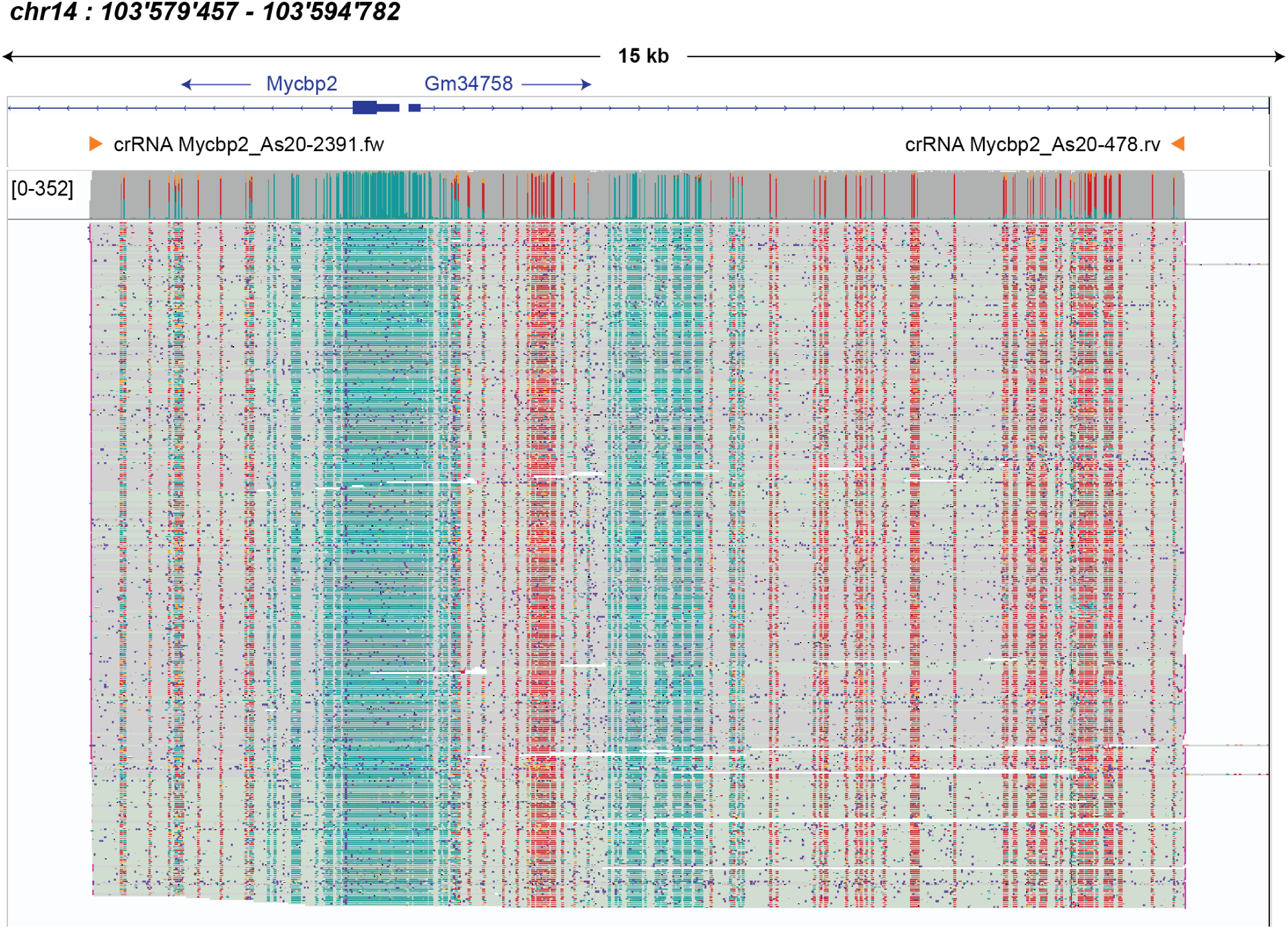
IGV-Snapshot of the reads aligned to the Myccb2 target region. Reads were obtained by simultaneously targeting 8 different loci (Supplementary Figures 7 to Supplementary Figure15, summarized in Figure 3). Forward reads are grey reverse reads light green, unmethylated (methylation probability <50%) CpGs are shown in teal, 5mC modifications are shown in red, and 5hmC modifications are shown in orange.

**Supplementary Figure 13:**
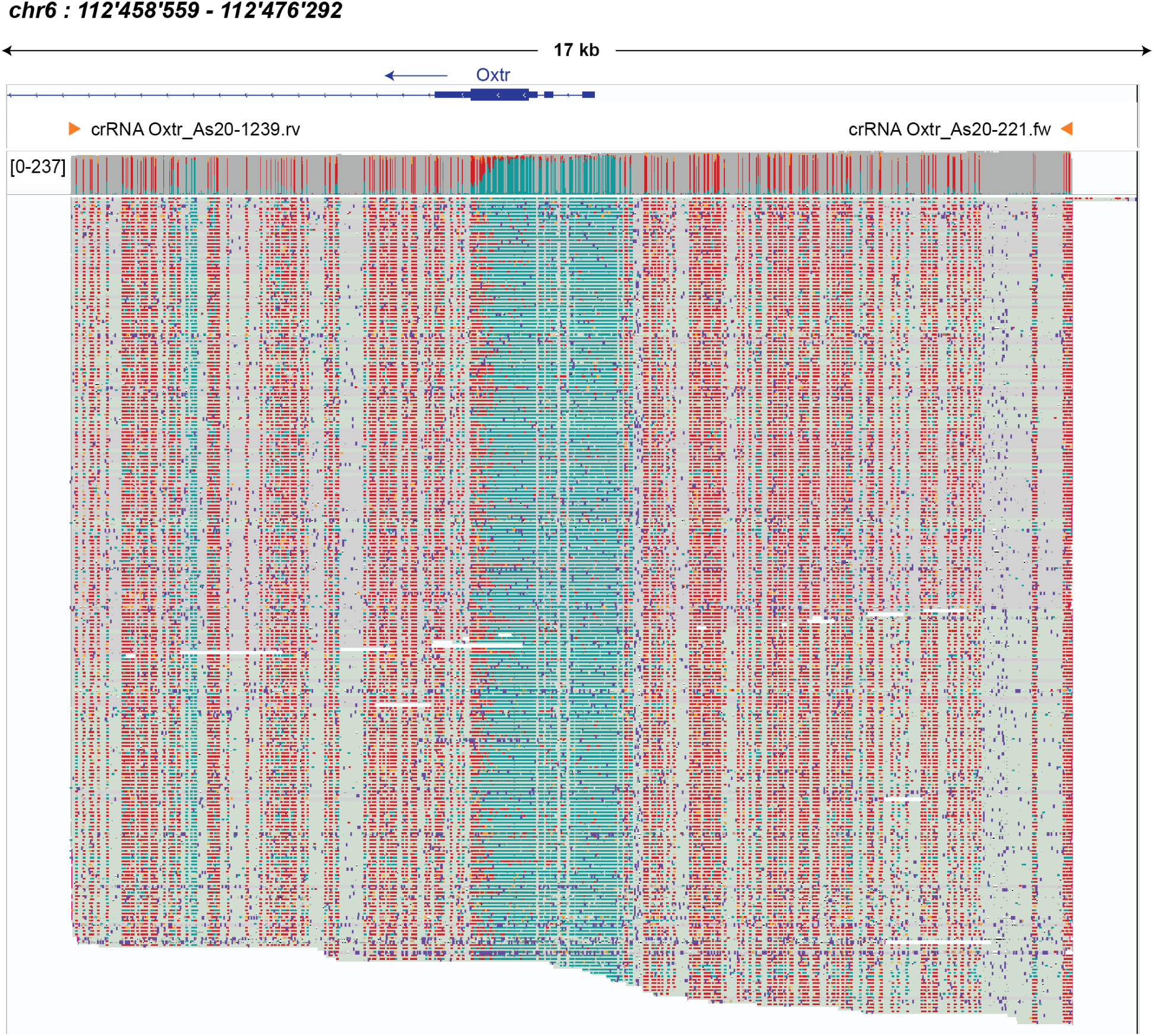
IGV-Snapshot of the reads aligned to the Oxtr target region. Reads were obtained by simultaneously targeting 8 different loci (Supplementary Figures 7 to Supplementary Figure15, summarized in Figure 3). Forward reads are grey reverse reads light green, unmethylated (methylation probability <50%) CpGs are shown in teal, 5mC modifications are shown in red, and 5hmC modifications are shown in orange.

**Supplementary Figure 14:**
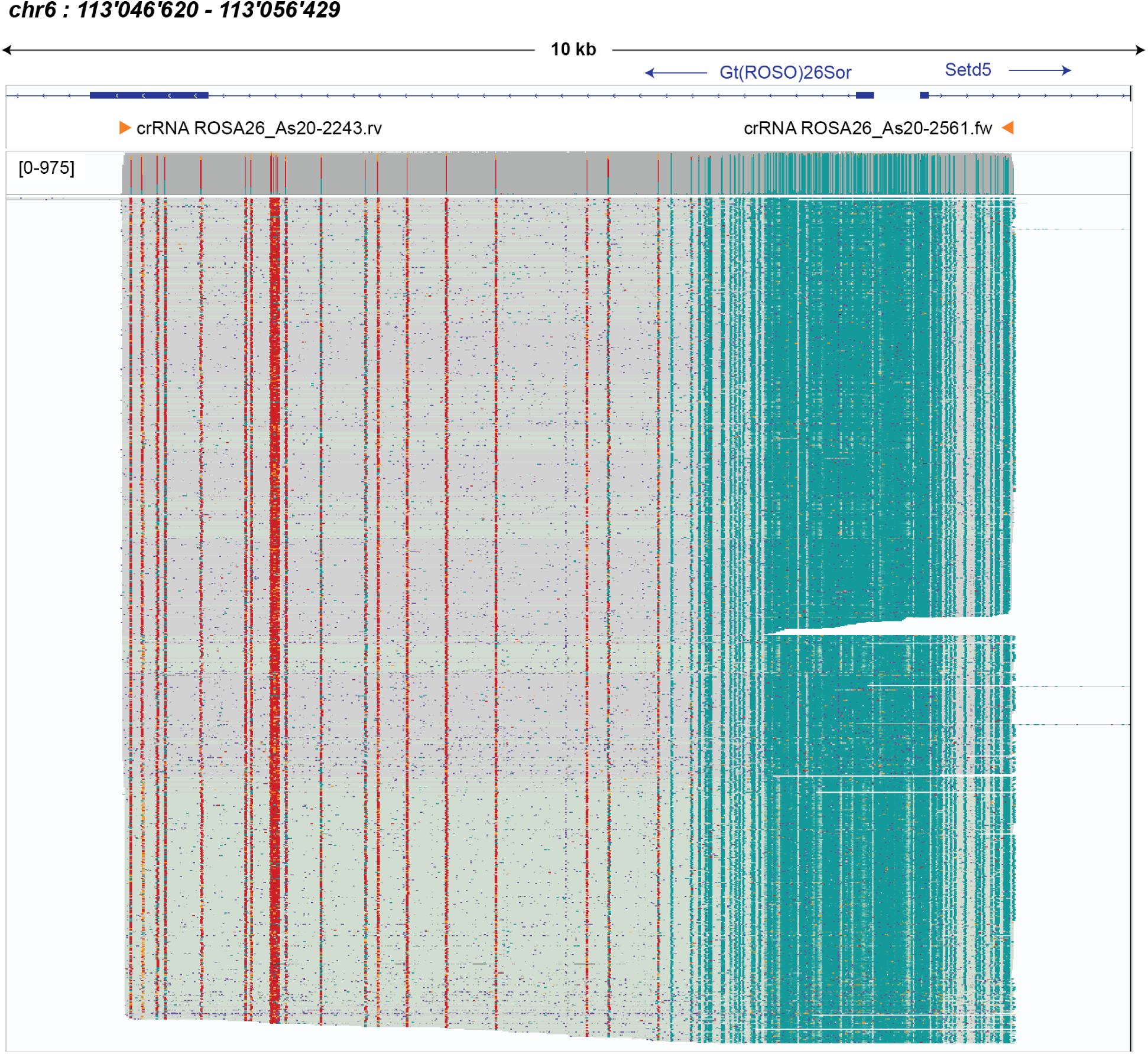
IGV-Snapshot of the reads aligned to the ROSA26 target region. Reads were obtained by simultaneously targeting 8 different loci (Supplementary Figures 7 to Supplementary Figure15, summarized in Figure 3). Forward reads are grey reverse reads light green, unmethylated (methylation probability <50%) CpGs are shown in teal, 5mC modifications are shown in red, and 5hmC modifications are shown in orange.

**Supplementary Figure 15:**
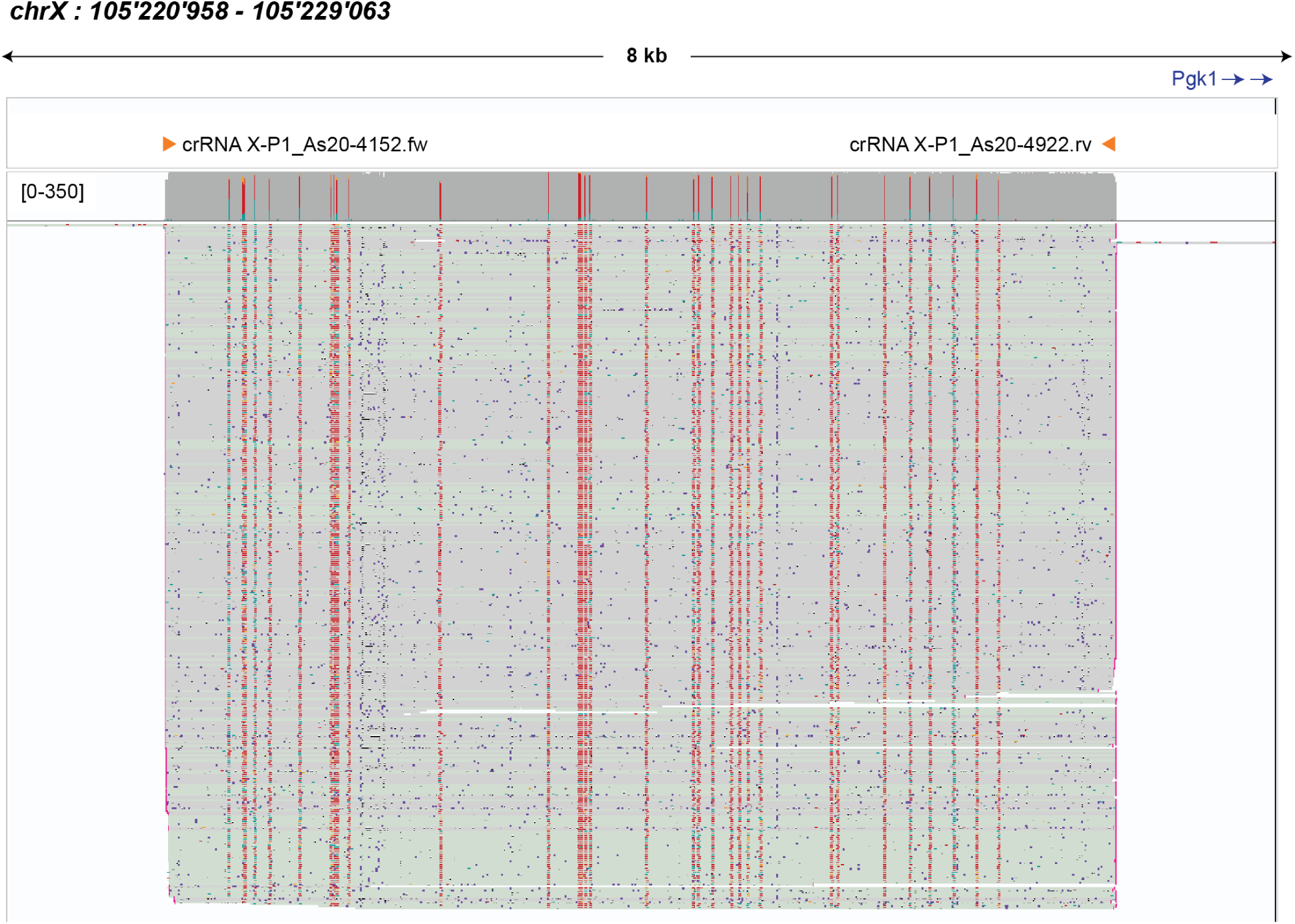
IGV-Snapshot of the reads aligned to the XP1 target region. Reads were obtained by simultaneously targeting 8 different loci (Supplementary Figures 7 to Supplementary Figure15, summarized in Figure 3). Forward reads are grey reverse reads light green, unmethylated (methylation probability <50%) CpGs are shown in teal, 5mC modifications are shown in red, and 5hmC modifications are shown in orange.

**Supplementary Figure 16:**
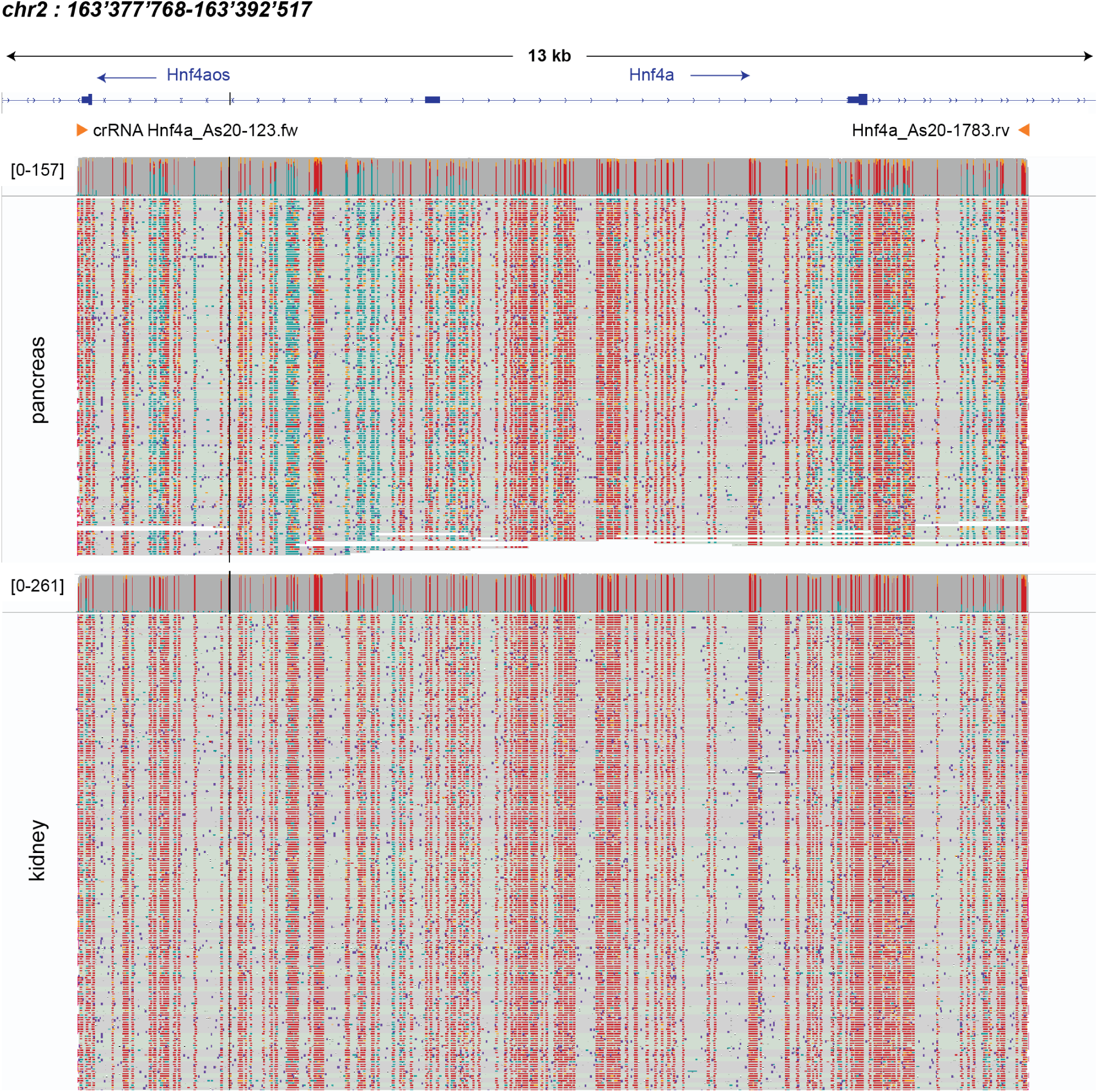
IGV-Snapshot of the reads from kidney and pancreas aligned to the Hnf4a target. Reads were obtained by barcoded multiplexing of 4 different tissues using CMT-nSeq (see also Figure 4). Forward reads are grey reverse reads light green, unmethylated (methylation probability <50%) CpGs are shown in teal, 5mC modifications are shown in red, and 5hmC modifications are shown in orange.

**Supplementary Figure 17:**
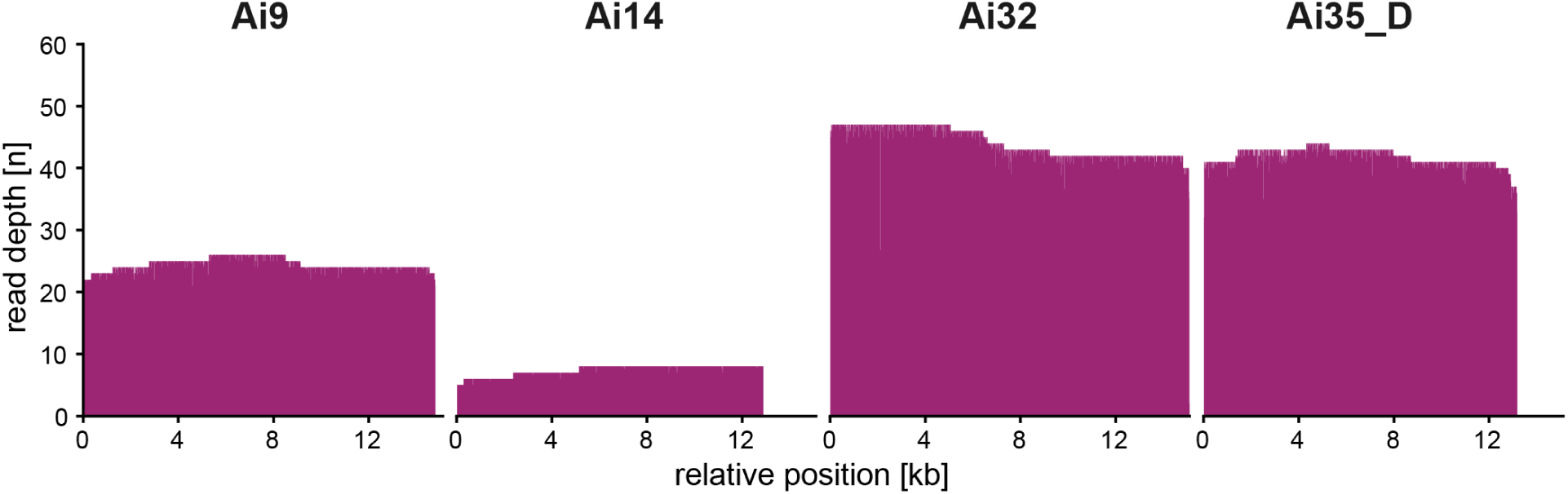
Coverage of targeted ROSA26 loci achieved using multiplexed size-enriched nCATS. The nCATs protocol was adapted to R10 and combined with fragment size enrichment on a 0.5% agarose gel, positions are relative to the PAM of the most upstream crRNA.

**SUPPLEMENTARY TABLE 1:**
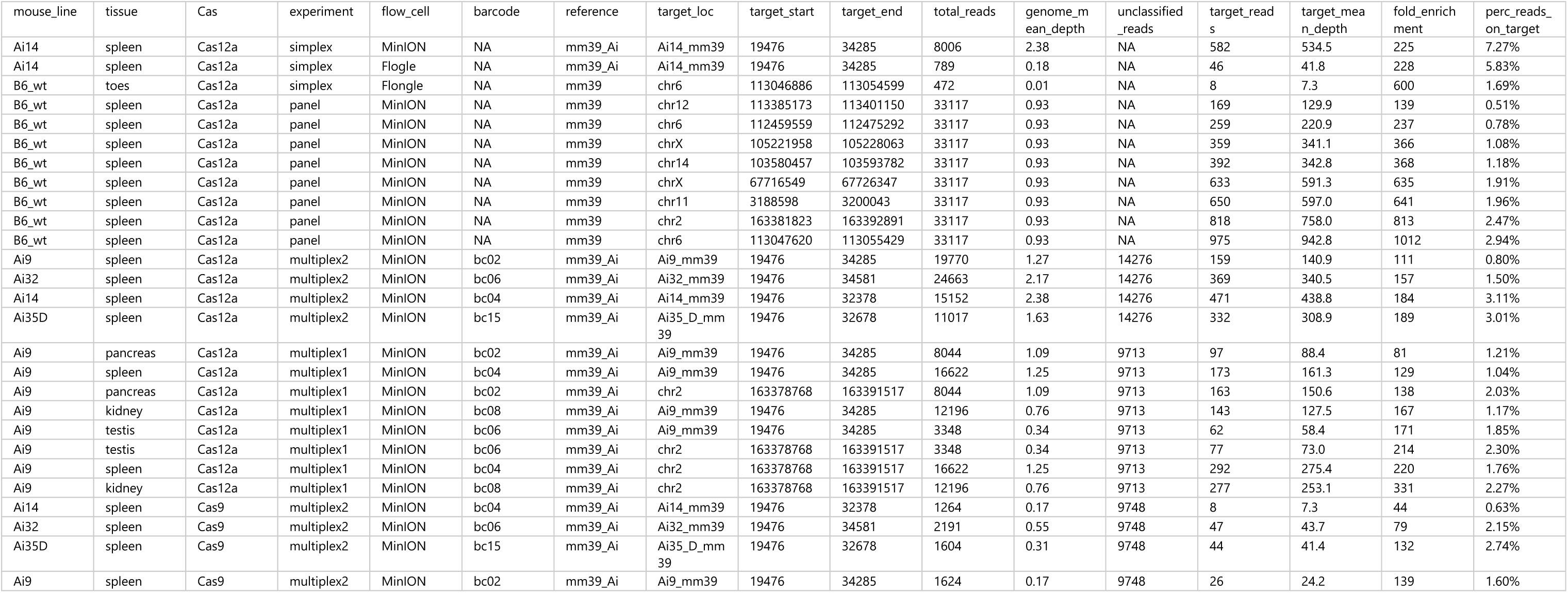
OVERVIEW ON READ STATISTICS FOR ALL TARGETED ONT SEQUENCING EXPERIMENTS.

**Supplementary Table 2:**
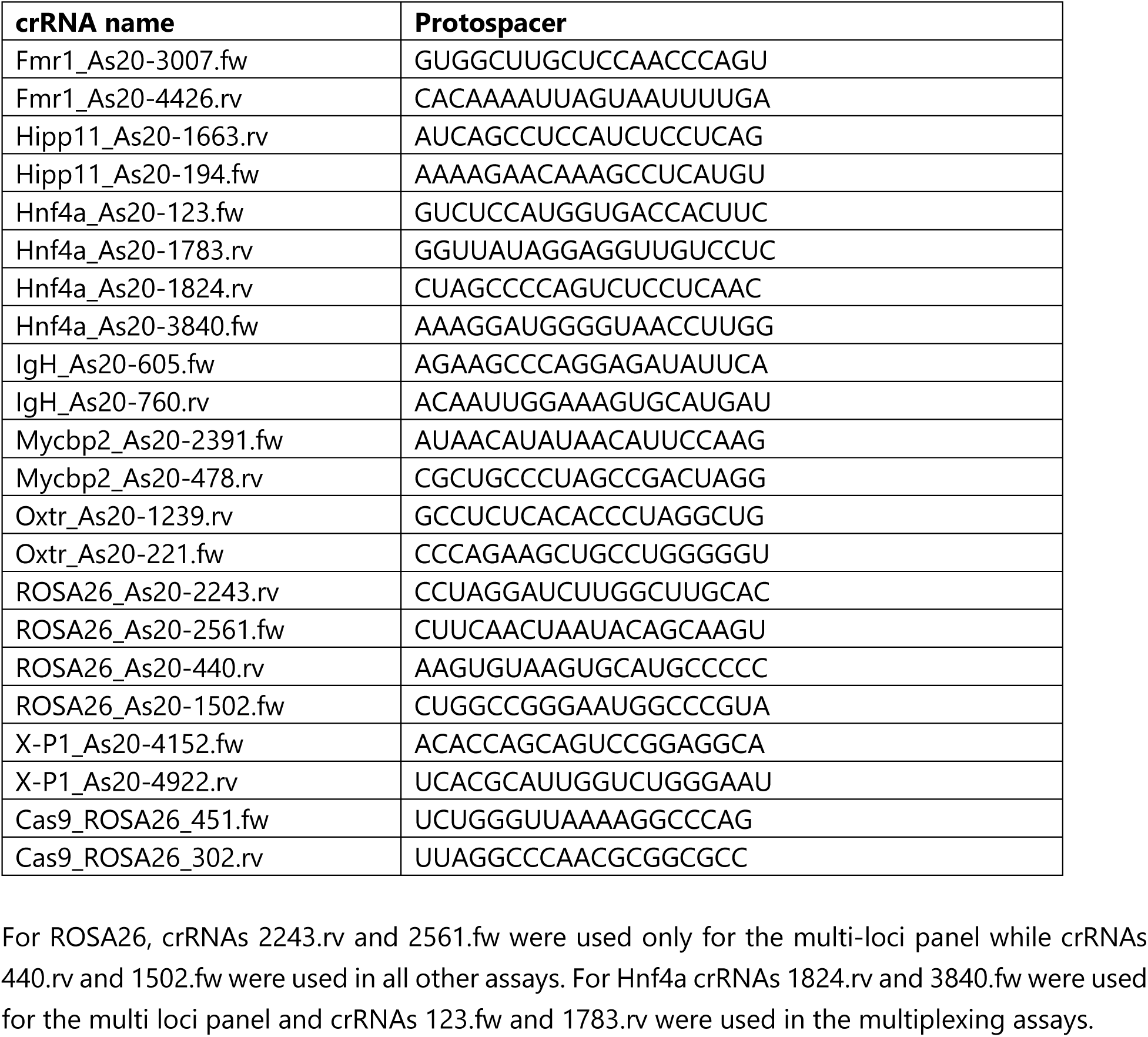
Protospacer sequences of crRNAs.

**Supplementary Table 3:**
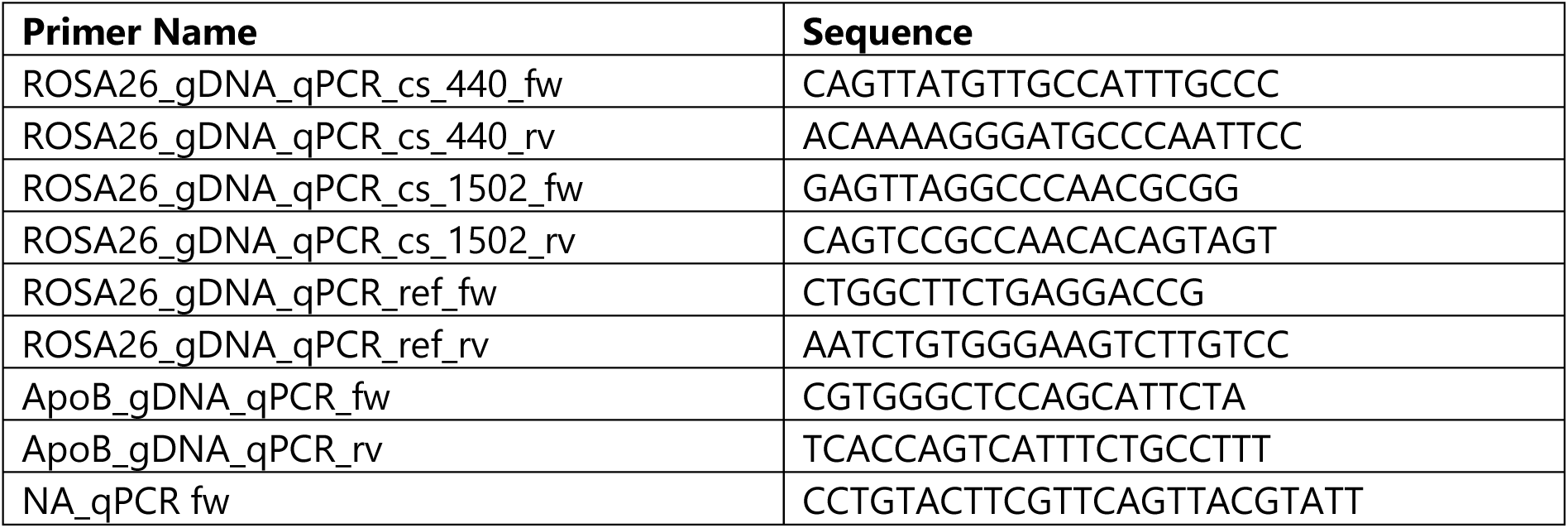
qPCR Primer sequences.

**Supplementary Table 4:**
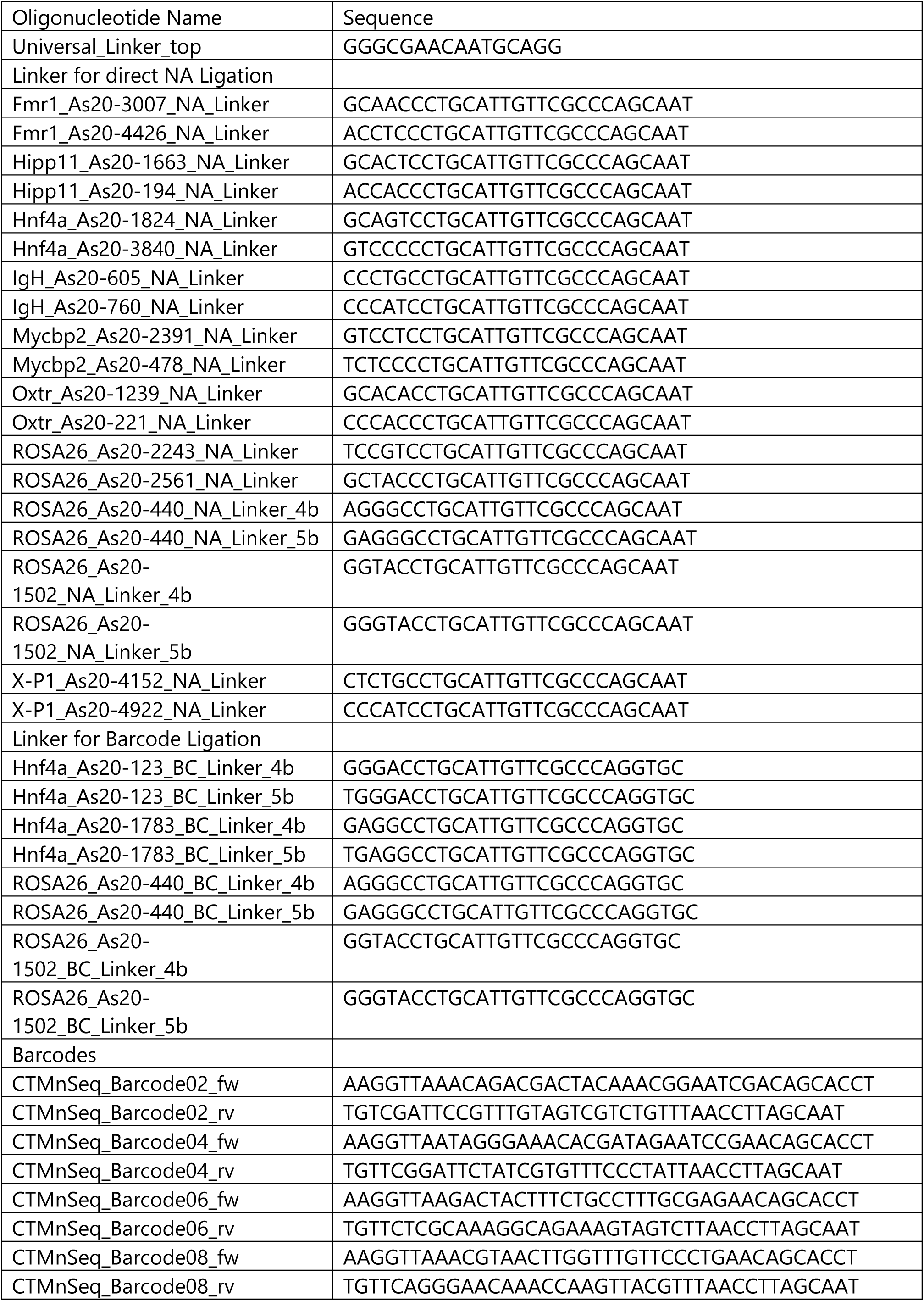
Sequences of DNA oligonucleotides used as linkers and custom barcodes.

**Supplementary Table 5:**
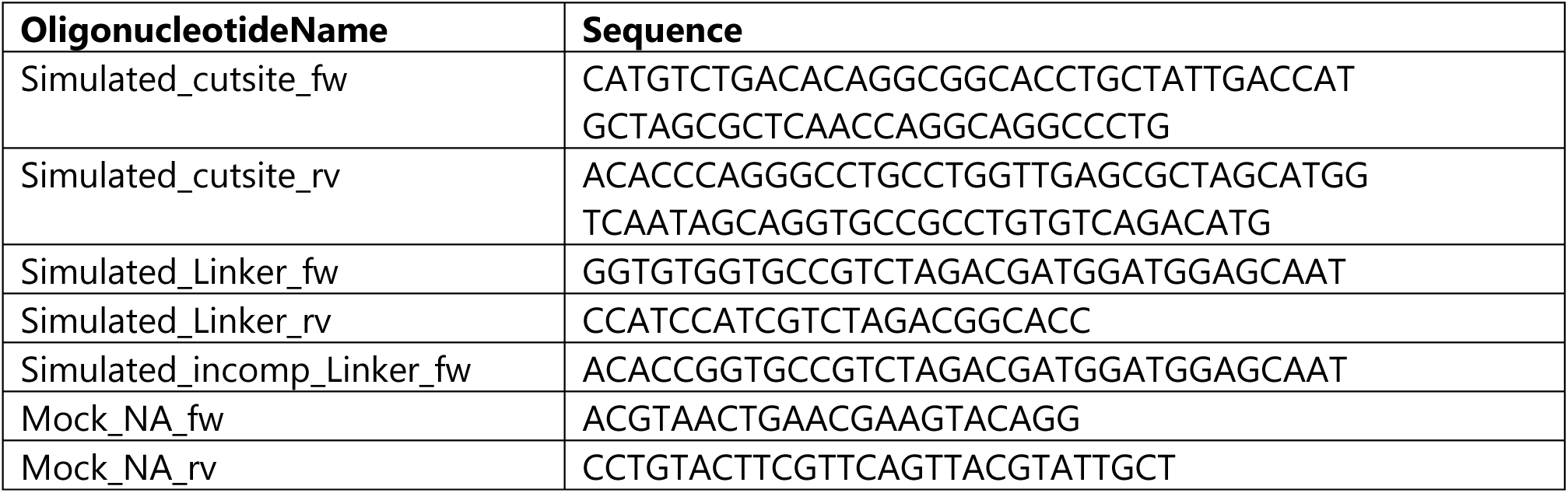
Oligonucleotides for generating simulated cut-site, linker and NA used for ligation tests.

